# Bayesian nonparametric analysis of residence times for protein-lipid interactions in Molecular Dynamics simulations

**DOI:** 10.1101/2024.11.07.622502

**Authors:** Ricky Sexton, Mohamadreza Fazel, Maxwell Schweiger, Steve Pressé, Oliver Beckstein

**Affiliations:** Department of Physics, Arizona State University, Tempe AZ, USA; Center for Biological Physics, Arizona State University, Tempe AZ, USA; School of Molecular Sciences, Arizona State University, Tempe AZ, USA; National Cancer Institute, National Institute of Health, Bethesda, MD 20892, USA

## Abstract

Molecular Dynamics (MD) simulations are a versatile tool to investigate the interactions of proteins within their environments, in particular of membrane proteins with the surrounding lipids. However, quantitative analysis of lipid-protein binding kinetics has remained challenging due to considerable noise and low frequency of long binding events, even in hundreds of microseconds of simulation data. Here we apply Bayesian nonparametrics to compute residue-resolved residence time distributions from MD trajectories. Such an analysis characterizes binding processes at different timescales (quantified by their kinetic off-rate) and assigns to each trajectory frame a probability of belonging to a specific process. In this way, we classify trajectory frames in an unsupervised manner and obtain, for example, different binding poses or molecular densities based on the timescale of the process. We demonstrate our approach by characterizing interactions of cholesterol with six different G-protein coupled receptors (A_2A_AR, *β*_2_AR, CB_1_R, CB_2_R, CCK_1_R, CCK_2_R) simulated with coarse-grained MD simulations with the MARTINI model. The nonparametric Bayesian analysis allows us to connect the coarse binding time series data to the underlying molecular picture and, thus, not only infers accurate binding kinetics with error distributions from MD simulations but also describes molecular events responsible for the broad range of kinetic rates.

## 1 Introduction

Membrane proteins exist in the plasma membrane of the cell and in the membranes of intra-cellular organelles to carry out functions such as ion and small molecule transport and signal transduction across the membrane. Although these proteins exist only in membranes, nearly one third of all proteins are membrane proteins.^1^ Intrinsic membrane proteins span the space of the hydrophobic lipid tails of the membrane^2^ and perform functions ranging from transport of ions and small molecules across the membrane to signal transduction with small-molecule agonists. Many membrane proteins are targets for drugs for treating a wide range of conditions, from fatigue^3^ to Parkinson’s disease.^4^ The local membrane environment is important for proper functioning of these proteins, as evidenced by the effect of lipid composition,^5–17^ mechanical membrane deformations,^18,19^ changes in membrane voltage,^20,21^ and even light,^22,23^ as well as the presence of activating ligands.^24,25^ Although some membrane protein-environment interactions such as the modulation of G protein-coupled receptor (GPCR) function by cholesterol are well known, the molecular mechanisms are not always well understood.

GPCRs are membrane proteins passing signals through the membrane by catalyzing the dissociation of intracellular G proteins^4,26–29^ in response to an extracellular binding event. The function of some GPCRs is known to be affected by cholesterol in the membrane, but the effects vary for different GPCRs.^5–17^ Signaling activity of the beta-2 adrenergic receptor (β_2_AR) increases when membrane cholesterol is depleted,^5^ but the oxytocin receptor (OTR) has the opposite response, becoming inactive in cholesterol depleted membranes.^6^ Homologous proteins such as the cannabinoid receptors (CB_1_R, CB_2_R) and cholecystokinin receptors (CCK_1_R, CCK_2_R) have shown differences in sensitivity to membrane cholesterol,^7–10,30,31^ but the mechanism of signal modulation by cholesterol is not well understood. The mechanism of cholesterol modulation has been attributed to both the membrane properties (such as stiffness) and specific interactions with cholesterol.

Molecular Dynamics (MD) simulations have been widely employed to directly probe the interactions of membrane proteins with lipids and to determine the specific interaction sites. Several packages have been developed to aid in the analysis of MD simulations for protein-lipid interactions.^32–38^ A key quantity to characterize the interaction of a protein residue with a small molecule such as a lipid, a drug, or solvent, is the residence time (τ), quantifying the time that an interaction persists until the small molecule unbinds. For a simple kinetic model of binding of the ligand X to the protein 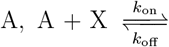, implying an exponential waiting time distribution, the inverse of the residence time is the off-rate, *k*_off_ = *τ*^−1^. In drug discovery, *τ* is a key quantity to characterize the strength of interaction between a small molecule and a protein.^39^ For easily accessible binding sites on protein surfaces, such as the ones for lipids interacting with membrane proteins, the on-rate *k*_on_ may be close to the diffusion limit and thus fairly similar for different binding sites so that *τ* would primarily distinguish these sites. In the general case, the on-rate together with the off-rate is required for the full kinetic characterization of binding but in this work we focus on the off-rate (or residence time) alone due to its relevance for protein-lipid interactions and for methodological simplicity because, unlike the residence time, the time until binding depends on the ligand concentration and thus requires careful and extensive sampling of the lipid environment surrounding a binding site. Previous studies computed the residence time in different ways, including taking the mean of all contact event times (when the protein residue lipid distance remains below a chosen cutoff distance),^40^ using autocorrelation functions of the time series of contacts,^40–42^ fitting the residence time survival function to an exponential distribution,^39,43^ computing directly from the residence time survival function,^40^ and using enhanced sampling methods.^44–49^ In this work we refer to the individual observed binding event times as the residence (or waiting) times and the timescale of the slowest unbinding process (discussed in detail below) as simply *τ*.

In prior work, a single exponential or two-term exponential mixture model for the waiting time distribution was least-squares fitted to the survival function computed from the observed waiting times to provide a point estimate for the slowest off-rate, taken to represent the binding process.^32,39,43,50^ However, in practice, the residence time distributions do not follow simple single or double exponential distributions. Firstly, the use of cutoffs in the contact analysis introduces fast artificial “flicker” events during which contacts are detected when the distance between protein residue and lipid fluctuates around the cutoff distance. More importantly, however, is that a simple binding criterion such as a distance to a single residue may not discriminate adequately between different molecular binding processes and thus lead to distributions that are mixtures of multiple un-known processes. In this case, such simple fitting approaches result in estimates for *τ* that poorly match the data in the long time-scale regime with inadequate estimates for the true error. (To further illustrate the shortcomings of the least square fitting approach, we provide a comparison of it to the method introduced in this work in the Supplementary Information in Section S1 and figure S1.)

Here we describe a method to analyze contact time series data under the hypothesis that they represent a mixture of *K* distinct binding processes and thus their waiting time distribution can be modeled as a *K*-term exponential mixture model (also known as the hyperexponential distribution)

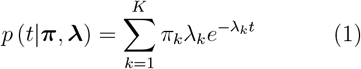

where *t* is a waiting time, **π** = {*π*_1_, …, *π*_*K*_} are the weights for the different components, and ***λ*** = {*λ*_1_, …, *λ*_*K*_} are the rates for the *K* processes. As we do not know the number of components *K* in advance, we take *K* to be very large (technically infinite) and allow Bayesian nonparametric inference^51^ to winnow down the number of mixture components contributing significantly to the data, along the associated per-residue residence times. Our method improves the estimates for 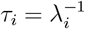 obtained from MD simulations and provides error estimates incorporating all available data.

Additionally, the Bayesian approach enables us to assign to each binding event a probability to have been sampled from one of the *K* processes, which in turn allows us to directly analyze the MD trajectory in terms of the different processes and in this way cluster the simulation by the inferred time scales. We call this novel approach *kinetic mapping*.

The paper is organized as follows: We first describe how waiting-time time series are obtained from MD simulations. We then derive the Bayesian nonparametric inference scheme and the resulting posterior distributions over the weights of each component, sampled numerically with a Gibbs sampler. We then obtain an estimate of the timescales of all relevant processes that contributed contacts at a specific residue. We validate our approach on a synthetic data set and describe how to perform analysis of MD trajectories to obtain different binding poses and ligand densities for processes with different off-rates. We apply our method to the problem of identifying specific cholesterol-GPCR interactions sampled in coarse grained (CG) MD simulations with the MARTINI force field^52–55^ as described in our earlier work.^50^ In particular, we simulated the *β*_2_AR and A_2A_ adenosine receptor (A_2A_AR), which both contain well-characterized cholesterol binding sites, and the pairs of homologous receptors CB_1_R/CB_2_R and CCK_1_R/CCK_2_R, for which closely related proteins are known to have very different sensitivity to cholesterol.

## 2 Methods and Theory

Our approach consists of the following steps, which will be described in more details below (figure 1): (1) We sample the relevant interactions. In our case we focus on cholesterol-GPCR interactions that are obtained from CG MARTINI MD simulations. (2) We generate a time series of contact times **t** = {*t*_1_, *t*_2_, … *t*_*n*_, …} between each lipid and each residue. (3) We use Bayesian nonparametric inference together with a numerical Gibbs sampler to obtain the posterior distributions of the model parameters of Eq. 1. (4) We post-process the samples to quantify individual exponential components of the model. (5) In particular, we compute rates for the individual components (*i*.*e*., for the individual binding processes) as the *maximum a posteriori* (MAP) estimate from the samples of each component. With the important caveat that in our scheme complete results are full *joint distributions* over all parameters, thus, wherever point-estimates are of interest, we choose to show the MAP estimate, since it is, by definition, the point-estimate that suits the data the best.^51^ (6) As an optional step, we map MD trajectory frames to components (an approach that we call *kinetic mapping*) and analyze the simulation in terms of the different time scale processes.

**Figure 1:**
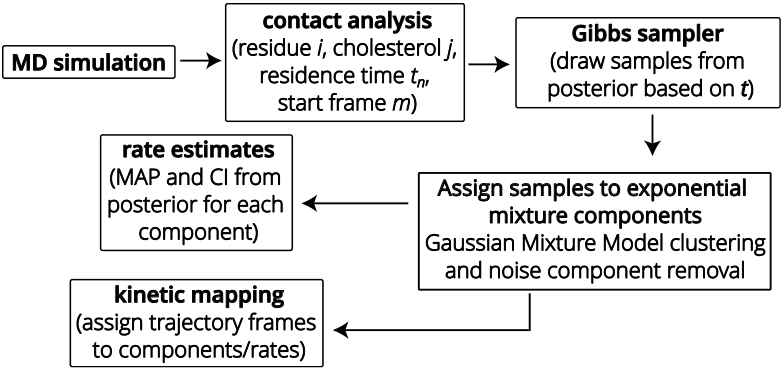
Workflow diagram for estimating off-rates for protein-cholesterol binding events from **MD simulations** by Bayesian nonparametric inference. Binding events are encoded into a time series of residence times (**contact analysis**). Samples from the posterior of the exponential mixture model Eq. (1) are obtained numerically with a **Gibbs sampler**. These samples are **assigned to the components of the exponential mixture model** using a clustering procedure and components insufficiently supported by data are removed as noise. **Rate estimates** are calculated for each component from the associated samples. Optionally, the original trajectory frames can be mapped back to components, thus clustering the trajectory by the inferred time scales (**kinetic mapping**).

### 2.1 System Preparation and Simulations

#### GPCR structures and models

We selected four GPCRs with resolved cholesterol molecules in experimental structures for testing our method: the adenosine receptor A_2A_ (A_2A_AR, structure with PDB ID 4eiy ^56^), the *β*_2_ adrenergic receptor (*β*_2_AR, PDB ID 2rh1^57^), and the cannabinoid receptors 1 (CB_1_R, PDB ID 5u09^58^) and 2 (CB_2_R, PDB ID 5zty^59^). All structure were obtained from the Protein Data-bank.^60^ All simulations for A_2A_AR and *β*_2_AR were taken from our previous work^50^ but for completeness we will include all relevant information here. Each of the structures had inserts on one of the loops to aid in crystallization, which were removed in the structure preparation and short linkers were added to fill the resulting gap in the structure using Modeller.^61,62^

We also include the two homologous cholecystokinin receptors (CCK_1_R and CCK_2_R), which we had analyzed previously,^50^ with additional MD simulations to better sample the very long residence times (> 1 *µ*s) that we observed in the previous CCK_1_R simulations.^50^ No structures had been available for either CCK_1_R or CCK_2_R at the time the simulation systems were created, so homology models were built based on the orexin receptors OX_1_R (PDB IDs 4zj8 and 4zjc^63^) and OX_2_R (PDB IDs 4s0v,^64^ 5wqc and 5ws3^65^) using Modeller:^61,62^ Inserts were removed from the structures and Jalview^66^ was used to perform a multiple sequence alignment with CCK_1_R and CCK_2_R, with sequences obtained from Uniprot.org^67^ (P32238 and P32239, respectively). Long loops were removed from the sequences and short linkers were used in their place. Using Modeller, twenty models for each protein were created and the structures with the smallest normalized DOPE scores^68^ were used as final models (−0.52 for CCK_1_R and −0.4 for CCK_2_R). More recently, crystal structures of CCK_1_R (PDB ID 7mbx^69^) and CCK_2_R (PDB ID 7f8w^70^) became available. In order to assess the quality of our models, we produced a structural alignment between the CCK_1_R and CCK_2_R models and the experimental structures with Chimera^71^ (figure S3 in Supplementary Information) and computed the backbone RMSD between the 271 and 257 matching residues with MDAnalysis, ^72^ which resulted in an RMSD of 2.83 Å for CCK_1_R, and 2.87 Å for CCK_2_R. Overall, the majority of the differences are located near the intracellular end of TM6 and in the extracellular region while the surfaces in contact with the membrane appear to be very similar between our models and the structures. Thus, even though our primary intention was to have results that could be consistently compared to our earlier work, ^50^ the good quality of the models, especially at the protein-lipid interface, suggests that results based on our models for CCK_1_R and CCK_2_R may be generally valid.

#### Coarse-grained MD simulations

The resulting models were coarse-grained in the MARTINI 2.2 force field^52–55^ and inserted into a 4:1 1-palmitoyl-2-oleoyl-sn-glycero-3-phosphocholine (POPC):cholesterol membrane using the CHARMM-GUI server.^73^ Elastic Network dynamic (ElNeDyn)^74^ secondary structure restraints were added to maintain the structure of the coarse-grained protein. MD simulations were performed with GROMACS 2019^75^ at 303.15 K and 1 bar with semiisotropic Parrinello-Rahman pressure coupling^76^ and stochastic velocity-rescale temperature coupling^77^ with a 20 fs integrator time step. Electrostatics were calculated with a reaction-field using a 1.1 nm cutoff and van der Waals interactions were computed with a 1.1 nm single cutoff, as described in our previous work.^50^ Some simulations of CCK_1_R and CCK_2_R were performed with GROMACS 2023 due to changes in computational environment, with further details such as system sizes and run lengths listed in Table 1.

**Table 1.** CG MD simulations of six GPCRs analyzed in this work. Some simulation data were taken from earlier work as indicated.

| Protein | Simulation length (μs) | #Particles | #Lipids | Box dimensions ($\text{\AA}$)(x,y,z) | GROMACS | Ref |
| --- | --- | --- | --- | --- | --- | --- |
| A<sub>2</sub>AAR | 109.9 | 10489 | 293 | (94.1, 94.1, 129.5) | 2019.2 | Geiger et al.<sup>50</sup> |
| β<sub>2</sub>AR | 121.5 | 9260 | 291 | (94.1, 94.1, 113.3) | 2019.2 | Geiger et al.<sup>50</sup> |
|  | 120.9 |  |  |  |  |  |
|  | 123.8 |  |  |  |  |  |
| CB<sub>1</sub>R | 100.6 | 8187 | 288 | (93.8, 93.8, 99.5) | 2019.2 | this work |
| CB<sub>2</sub>R | 101.2 | 8197 | 287 | (93.3, 93.3, 101.0) | 2019.2 | this work |
| CCK<sub>1</sub>R | 100.3 | 9102 | 293 | (95.6, 95.6, 107.6) | 2019.2 | Geiger et al.<sup>50</sup> |
|  | 66.3 |  |  |  | 2023.4 | this work |
|  | 67.6 |  |  |  |  |  |
|  | 66.9 |  |  |  |  |  |
| CCK<sub>2</sub>R | 103.3 | 8909 | 292 | (95.5, 95.5, 105.8) | 2019.2 | Geiger et al.<sup>50</sup> |
|  | 58.1 |  |  |  | 2023.4 | this work |
|  | 56.8 |  |  |  |  |  |
|  | 55.1 |  |  |  |  |  |
|  | 54.3 |  |  |  |  |  |

### 2.2 Residence Time Analysis

The MD simulations sample coordinates for all particles in the system at regular time steps, to which we refer as “trajectory frames”. We quantify protein-lipid interactions with a simple contact analysis. A protein residue *i* and a lipid molecule *j* were considered in contact whenever any lipid CG particle bead’s position 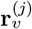 (the coarse-grained equivalent of an atom) was within a cutoff distance *d* of any of the residue bead positions 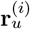, *i*.*e*., a contact at trajectory frame *n* is recorded if 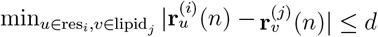.

To determine the proper cutoff, the minimum distance between protein and lipids was calculated for each frame for each residue and histogrammed, three of which are shown in figure S4 in the Supplementary Information. The histograms consistently showed minima above 6 Å and below 8 Å, so a cutoff of 7 Å was chosen, similar to (but slightly larger than) the value of 6.3 Å used in a previous study.^78^ The time series of the distance to each lipid was calculated for several residues over the trajectory, which showed that a cutoff of 7 Å typically included fluctuations from the closest position without including long binding events from nearby residues. A representative lipid-residue distance time-series is plotted with a dashed horizontal line drawn at 7 Å in figure 2, along with cholesterol binding poses corresponding to the contacts depicted in the time-series. For a chosen cutoff, a residence time series was collected for each residue, where a contact event was recorded with its start time *s*_*n*_ (i.e., the index number of the frame in the trajectory), duration *t*_*n*_, and the index of the lipid forming the contact *j*. All lipids of the same chemical composition that interact with the specific protein residue contribute to the same time series (figure 2).

**Figure 2:**
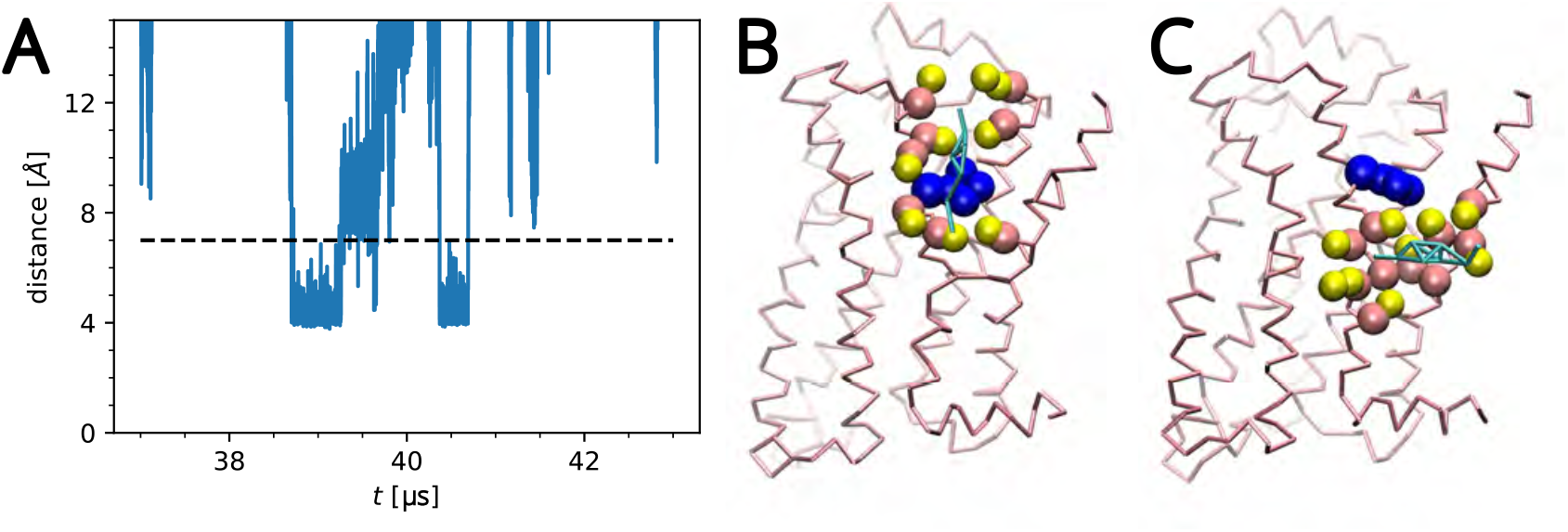
Representative cutoff example: (A) Time-series of a W313-cholesterol distance for a single cholesterol. (B) Cholesterol (cyan) bound to W313 (blue) at time 39 *µ*s. Residues within 7 Å are shown in the VdW representation. (C) The same cholesterol at 39.5 µs unbound from W313 but bound to nearby residues.

We assessed the sensitivity of our approach to the choice of cutoff for the *β*_2_AR data set. We calculated the maximum residence time τ for cutoffs 6 Å, 7 Å, and 8 Å as shown in figure S2 and discussed in more detail in Section S3 in the Supplementary Information. The residence time *τ* increased with the length of the cutoff but generally the same specific residues were highlighted with longer than average residence times. The difference in τ between 6 Å and Å was more pronounced than between 7 Å and 8 Å, with the 8-Å results being the least precise ones as indicated by the largest confidence intervals. We therefore concluded that results are qualitatively fairly insensitive to the choice of cutoff in the interval that is reasonable for coarse-grained MD and that a cutoff of 7 Å provided a good balance between specificity and capturing longer binding events.

We directly used time series of residence times to infer kinetic parameters of binding. However, in order to summarize the data in a convenient form, we also used the normalized histogram of the residence times as an estimate for the underlying probability density of waiting times, *p*(*t*). The closely related survival function 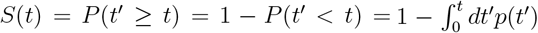 measures the probability to observe a binding event of at least length *t* and can be directly estimated from the data as an average over Heaviside functions of the waiting times *Ŝ*(*t*) = ⟨Θ(*t* − *t*_*n*_)⟩ or from the cumulative sum of the probability density. In a semi-log plot, the survival function shows a rapid drop with a slower decay at longer times, indicating that most observed contact events are short and that it is comparatively rare to observe long binding events (figure 3). The current state-of-the-art has consisted of fitting a two component exponential mixture to such survival functions, using a least squares regression^32^ (which can be improved upon by using relative weights^50^). However, the choice of only two terms in the exponential mixture is not justified and in fact it is not clear how many exponential components should be included, thus motivating the general *K*-term exponential mixture model in Eq. (1).

**Figure 3:**
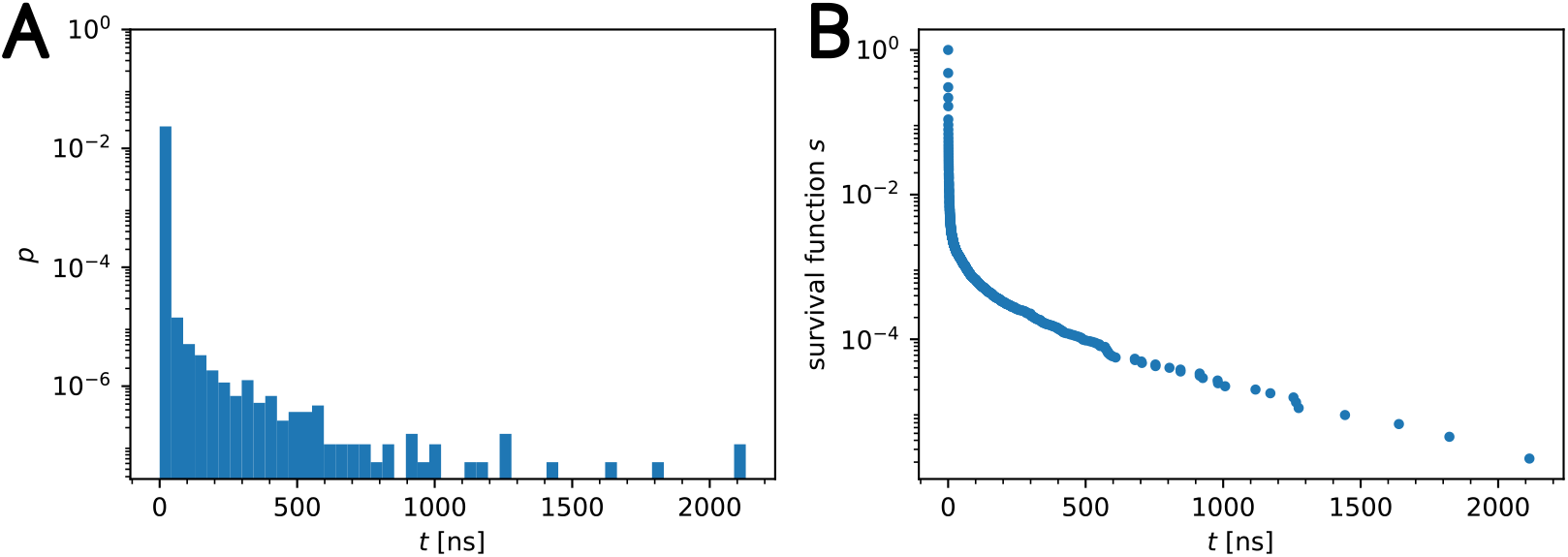
Representative data set (*β*_2_AR W313): Contacts between cholesterol and W313 of *β*_2_AR were collected using a 7 Å cutoff. (A) Normalized histogram (probability density) of waiting times *p*(*t*) and (B) survival function *S*(*t*).

### 2.3 Bayesian Inference

#### Model

Contacts formed between a lipid and a protein residue can vary in binding pose, partial contacts with different parts of the lipid may occur, while very brief, weak contacts may either be genuine or an artifact of using a simple cutoff criterion. We consider these different binding modalities as different binding processes or components in our model. We model each component *k* with a rate *λ*_*k*_ and an exponential waiting time distribution *λ*_*k*_ exp(−*λ*_*k*_*t*); thus, the full waiting time distribution for *K* components is a *K*-term exponential mixture model

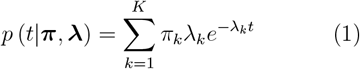

where *K* is the *a priori* assumed maximum number of mixture components, *t* is the observed residence time, and we want to determine the parameters of our model, namely, the weights ***π*** = {*π*_1_, …, *π*_*K*_} (which, when compared to a threshold probability as described in Section 2.5, determine whether each of the *K* possible components is indicated by the data) and the rates ***λ*** = {*λ*_1_, …, *λ*_*K*_}. As we want to calculate the distributions of our parameters for accurate error estimates and because the number of the *K* processes in the exponential mixture that are indicated by the data is not known *a priori*, we turn to Bayesian nonparametric inference.

#### Bayesian nonparametric inference

Bayes’ theorem describes the probability of an event based on prior knowledge updated using the likelihood,^51^

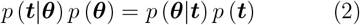

where the data ***t*** are the set of observed residence times (***t*** = {*t*_1_, …, *t*_Ω_}) and ***θ*** is the set of parameters (***π, λ***). Eq. (2) relates the likelihood of a set of residence times being observed given a set of parameters (*p*(***t***|***θ***)), to the (posterior) probability for a set of parameters given the data set (*p*(***θ***|***t***)). Since the objective is to determine the full posterior distribution over parameters given the observed data, Eq. (2) is rewritten as

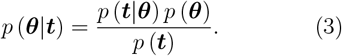

Here *p*(***θ***) is the prior, and *p*(***t***) = ∫*p*(***t***|***θ***) *p*(***θ***) *d****θ*** is just a normalizing factor, termed the evidence, that can be calculated later if needed. The full treatment of Bayes’ theorem with all parameters along with derivations of the conditional posteriors is located in the Supplementary Information.

In terms of selecting priors, a prior must be chosen so as to have support over all possible parameter values over which the posterior may exist. For computational reasons alone, it is often convenient to impose an additional constraint and seek a prior resulting in a conjugate prior-likelihood pair, *i*.*e*., guaranteeing that the prior and the posterior have the same form.^51^ This is often a useful choice since the posterior can then be determined in closed form and sampled directly. In fact, wherever possible, we keep priors with closed form posteriors in mind because direct sampling significantly improves the efficiency—both the speed and effectiveness—of each MCMC iteration. By contrast, more physically significant priors would require expensive computations—for example in Metropolis-Hastings, repeatedly computing expensive accept/reject criteria. Furthermore, the choice of priors becomes less important as the number of data points increases and in our case, the number of data points in a set are generally large (∼10^5^). Their overall contribution to the posterior diminishes and the shape of the posterior is ultimately dictated by the likelihood appearing in Eq. (3), as will be shown to be the case here.

In the present model, the conjugate prior on the weights (**π**) is a Dirichlet distribution

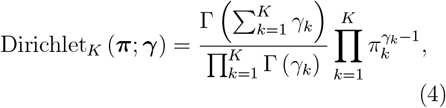

and the prior on the rates (**λ**) is a gamma distribution

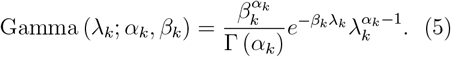

The distributions of the parameters depend on their own (hyper) parameters ***α, β, γ***. Hyperparameters are chosen such that the prior has a broad, diffuse mode. We specify our choice of hyperparameters after a full discussion of our model below.

The protein-lipid contacts for a given site can vary in several ways, from different orientations of the lipid to partial contacts on different parts of the lipid. The residence time distribution is sampled from a combination of these processes, but the amount each binding mode contributes to the total residence time distribution is unknown. To model our lack of knowledge we introduce a latent variable—the indicator **z** = (*z*_1_, …, *z*_*n*_, … *z*_Ω_)—to indicate which of the K components gave rise to each of the Ω data points, *i*.*e*., the indicator is a mapping from data points with index 1 ≤ *n* ≤ Ω to *K* components, *z*: *n* ↦ *z*_*n*_, *z*_*n*_ ∈ {1, …, *K*}. As the prior for the indicator’s posterior distribution we chose a categorical distribution,

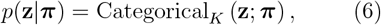

which describes drawing of samples from a set of categories based on an associated set of probabilities,^51^ namely the weights **π**. As shown in Section S2 of the Supplementary Information, the conditional posterior of the indicator is then

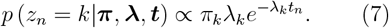

The use of the indicator allows for the determination of a closed form for the posterior distributions. Inserting the likelihood and priors into Eq. (3) as well as using the indicator and simplifying (as shown in Section S2 in the Supplementary Information) yields the marginal posterior distributions for the weights

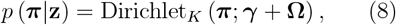

and rates

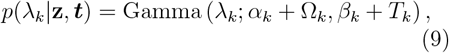

where **Ω** indicates how many data points are associated with each component of the model (**Ω** = {Ω_1_, …, Ω_*K*_} and 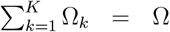). ***T*** = {*T*_1_, …, *T*_*k*_, …, *T*_*K*_} are the total residence times attributed to each component *k* with 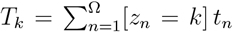 where the Iverson brackets select those values that belong to component *k*.

As can be seen in Eqs. (8) and (9), the hyperparameters add pseudocounts to the number of data points and total residence time for each component. Since both Ω and the *T*_*k*_ are typically large, a choice of hyperparameters on the order of one will suffice. In the present work the values for ***α*** and ***β*** were set to 1 and 3, respectively, so as not to exclude states with weights and rates near zero, since the quantity of interest is the timescale of the slowest process. The values for ***γ*** were all set to 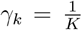 for every component so as to ensure a result independent of *K*;^51^ this choice allows us to choose a large but finite *K* for the practical implementation of our Bayesian nonparametric framework.

It is not possible to write an analytical expression for the posterior because the number of data points attributed to each component **Ω** and the total associated residence times **T** are not known and only implicitly described by the indicator **z**. We therefore resort to numerically sampling from the posterior. The classification of data points to each component in the model (Eq. (7)) and the conditional posteriors (Eqs. (8) and (9)) were sampled using Markov Chain Monte Carlo (MCMC). Since the posterior is naturally multivariate and the parameters can be grouped by weights and rates, a Gibbs sampler was used.

### 2.4 Gibbs Sampler

The Gibbs sampling scheme is an iterative algorithm that produces one sample of the weights and rates (**π, λ**) for each iteration, where the samples are distributed according to the conditional posteriors Eqs. (8) and (9) (Algorithm 1). For the first step, the weights and rates must be initialized but the exact values are irrelevant as long as feasible values are chosen.^51^ From these starting values, the Gibbs sampler samples the indicator **z** from a categorical distribution with probabilities given by Eq. 7 (using the known **π** and **λ** as well as the data) and then proceeds to calculate **Ω** and ***T*** from the data. At this stage, the weights **π** can be resampled according to Eq. (8). With the updated weights, all *K* rates λ_*k*_ are sampled according to Eq. (9) and a new iteration with fully updated parameters is started. As the random number generator we used `numpy.random.Generator` in NumPy^79^ with default settings and drew random samples with the `multinomial()`, `dirichlet()`, and `gamma()` distribution samplers using the instantaneous parameters according to equations (7)–(9).

As the initial values almost certainly have very low likelihood, the sampling process initially requires a number of iterations before convergence to the target distribution. This so-called burn-in phase is later removed from the resulting Markov chain. Furthermore, the resulting Markov chain generates dependent samples so in order to generate nearly independent, identically distributed (iid) samples from the target posteriors, the Markov chain is sub-sampled (“thinned”) to remove correlations. ^51^ The first 10 000 samples were removed as the burn-in phase and every 100th iteration of the Gibbs sampler was saved to thin the samples. Our results were insensitive to the exact values used for burn-in and thinning, as shown in figure S5 in Supplementary Information where burn-in was varied between 5 000 and 20 000 samples and figure S6 for thinning parameters ranging from 50 to 200.

#### Algorithm 1

Exp.mixt.model Gibbs sampler

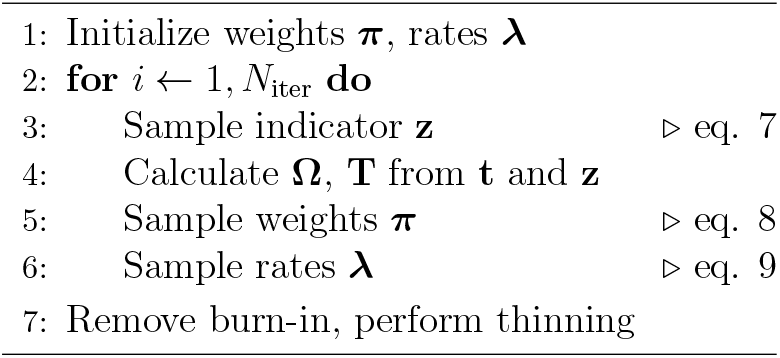

The total number of iterations of the Gibbs sampler (*N*_iter_) was set to 110 000 which, after burn-in removal and thinning, resulted in 1000 samples from the target posterior, which we considered iid. The samples were then post-processed and clustered to determine which samples belonged to which component of the exponential mixture model (see details in the next section) and to determine, for each cluster, the *maximum a posteriori* (MAP) estimate, *i*.*e*., the value of the parameters maximizing the posterior, and the 95% confidence interval of the samples for the error estimates.

As a practical matter, in order to run Algorithm 1 we may select a finite value for the number of mixture components *K*. We assume that only a small number of components *K*^′^ actually contribute to the total process. Thus, thanks to our choice of the *γ* hyperparameter, choosing *K* ≫ *K*^′^ would be essentially equivalent to choosing *K* = +∞.^51^ We ran the Gibbs sampler for *K* = 15, 20, and 50 on our representative test data set (“*β*_2_AR W313”, as described in Results). As figure 4 together with figures S7 and S8 and Table S1 in the Supplementary Information show, our results were very similar for these three values of *K* with only five components contributing per the criterion described in 2.5, demonstrating that our results were insensitive to this upper bound, provided it is set sufficiently high. We therefore chose *K* = 15 for all of our calculations.

**Figure 4:**
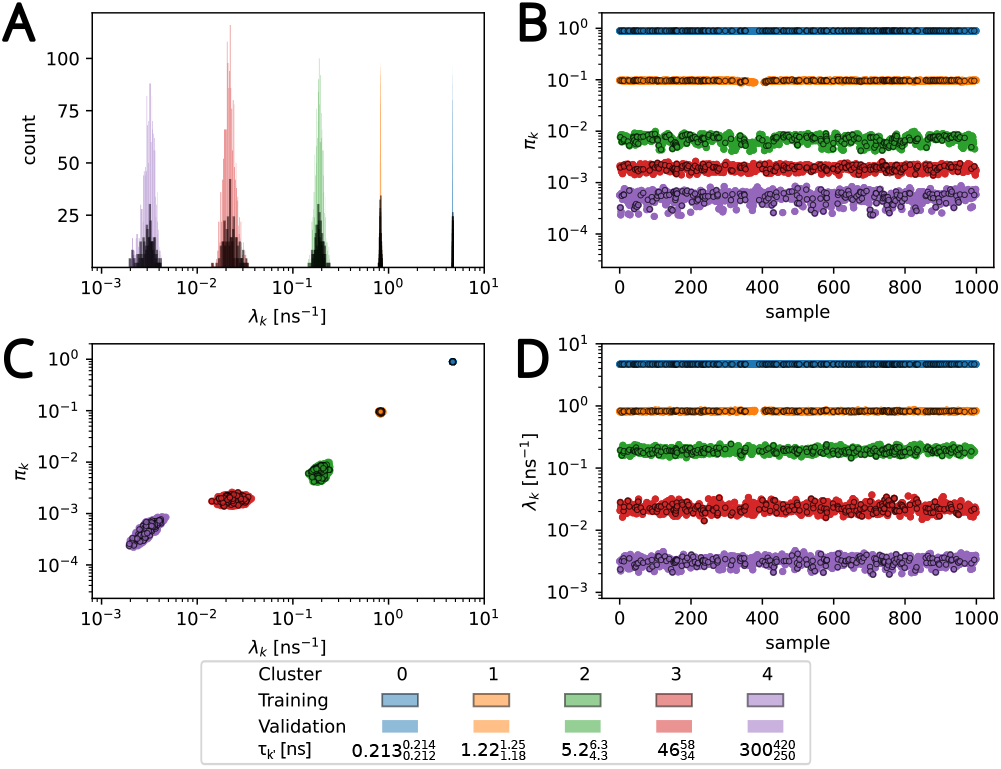
Weights and rates for the representative data set *β*_2_AR W313 from Bayesian nonparametric inference. (A) Histograms of rates λ_*k*_*′* (variable bin size), (B) weight π_*k*_*′* vs sample, (C) weight vs rate, and (D) rate vs sample for thinned Markov chains. Samples were clustered with a Gaussian mixture model (GMM) to correct for label-switching as described in Methods. Clusters *k*^′^ are colored and labeled in order of decreasing rate. Samples in the ‘training’ set were used to train the GMM and samples labeled ‘validation’ were fit to the trained model. Noise clusters were removed as described in Methods. Figure S12 in Supplementary Information depicts the same data with noise clusters included.

### 2.5 Label-switching correction

As the posterior at hand is degenerate with respect to (that is, invariant under changes to) the particular labeling of each mixture component,^51^ the label for a given state is not fixed. That is, in each independent MCMC sample, mixture components with the same physical role in the model are randomly assigned different indices. Counterintuitively, since this phenomenon, termed “label-switching” in the MCMC literature, replicates our posterior’s intrinsic degeneracy, observing label switching signals the physical soundness of our model; *i*.*e*. it indicates that the model on paper matches the model represented in code. However, practically, since meaningful indices are helpful to interpret our results, we will outline how we obtained physically interpretable labels here.

First, we must determine which mixture components from each MCMC sample are reinforced by the data, as using a Dirichlet prior on the weights naturally sets a non-zero *a priori* weight on all components. We achieve this with a simple cut-off of 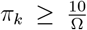 on each sample’s weights, removing samples with an expected number of associated contact events less than 10.

To determine which samples belong to each state, we followed Fazel et al. ^80^,^81^ and partitioned the samples into *K*^′^ clusters in parameter-space, each cluster corresponding to one component of the exponential mixture model Eq. 1. First, all samples with *K*^′^ weights, with *K*^′^ the *a posteriori* mode number, were used to train the `GaussianMixture` (GMM, Gaussian Mixture Model) estimator from scikit-learn^82^ in (log *λ*_*k*_, log *π*_*k*_) space. Then, this trained *K*^′^-clusters GMM estimator was used to partition the remaining samples into *K*^′^ clusters.

The cluster assignment mapped the indicator 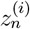 of a sample *i* for the data point *t*_*n*_ from the initial K components to the final *K*^′^ clusters, which resulted in a many-to-one mapping 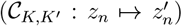 where several sampled components could ultimately be associated with a single cluster.

Rate estimates λ_*k*_*′* for each component *k*^′^ were determined from the maxima in the marginal posterior obtained from samples in cluster *k*^′^ along with 95% confidence intervals, *i*.*e*., to interpret our data we marginalized the full joint distribution over the weights and conditioned it on the cluster. The rate distribution was numerically transformed to a waiting time distribution by histogramming the reciprocals of the rate samples 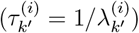. The maximum and 95% confidence interval of this distribution provided the MAP estimate and error for *τ*_*k*_*′*.

### 2.6 Noise cluster removal

The aforementioned clustering procedure produced sharply defined clusters together with diffuse clusters. We interpreted these diffuse clusters as noise, due to insufficient evidence in the data to confidently assign a data point *t*_*n*_ to a component of the model. We sought to identify and then exclude such *noise clusters* in order to automate analysis of large numbers of datasets, *e*.*g*., binding data for hundreds of residues. We identified noise clusters as follows: the marginal posterior probabilities for each data point *t*_*n*_ to belong to each of the *K*^′^ clusters were numerically obtained by histogramming the mapped cluster indicators 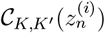 over all *w* samples,

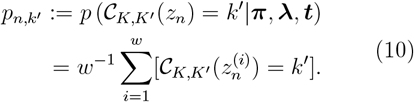

For example, a value of *p*_*n,k*_*′* = 0.6 indicates that waiting time data point *t*_*n*_ is associated with the model component *k*^′^ with probability 0.6 and its probability to belong to any other cluster except *k*^′^ is 0.4. Noise clusters were identified as those clusters *k*^′^ for which the (remapped) marginal posterior of the indicator *p*_*n,k*_*′* (Eq. 10) never exceeded 0.4 for *any* data *t*_*n*_, *i*.*e*., the probability for any observed waiting time to have belonged to the noise cluster would have been less than 0.4. The value of 0.4 was chosen heuristically as it robustly distinguished between ground truth components and noise for the synthetic data set.

The final estimate for the timescale of the slowest process was computed from the cluster with the slowest rate (or longest waiting time) after noise cluster removal, *i*.*e*., *τ* = max_*k*_*′ τ*_*k*_*′*.

### 2.7 Kinetic mapping

Since each data point *t*_*n*_ is associated with a specific binding event (protein residue number *i*, lipid number *j*, start time *s*_*n*_, duration *t*_*n*_) we can directly associate *t*_*n*_ with a trajectory segment starting at *s*_*n*_ and ending at *s*_*n*_ + *t*_*n*_. This *kinetic mapping* via the indicator (and the cluster mapping) associates the trajectory segment with the posterior probability distribution *p*_*n,k*_*′* = *p* (𝒞_*K,K*_*′* (*z*_*n*_) = *k*^′^ | ***π, λ, t***) (Eq. 10) for belonging to each of the *K*^′^ clusters. The segment can now be assigned to the cluster with the highest membership probability for *t*_*n*_ and in this way the trajectory is clustered by the timescales *τ*_*k*_*′* associated with the *K*^′^ clusters. Any standard trajectory analysis of molecular properties can now be performed over the collection of trajectory slices associated with a specific cluster and thus molecular properties of the specific process can be quantified. Alternatively, we may analyze the full trajectory but calculate properties of interest for each cluster *k*^′^ separately as weighted averages over the Ω trajectory segments with weights *p*_*n,k*_*′* for 1 ≤ *n* ≤ Ω.

### 2.8 Weighted Densities

For visualization with VMD^83^ and further analysis, we created trajectories representing the kinetic mapping for each residue of interest. Such trajectories only contained the protein and a single cholesterol molecule for all frames where cholesterol was bound to the residue of interest; note that this molecule may have had different identities (residue numbers) in the original trajectory but we treat them as indistinguishable because they have the same chemical identity. The probabilities that the binding event belonged to each cluster *p*_*n,k*_*′* (Eq. 10) were then associated with the appropriate frames.

Weighted spatial densities *ρ*_*k*′_ (**r**) were computed using a modified version of the `DensityAnalysis` analysis class in MDAnalysis.^72^ `DensityAnalysis` computes densities by histogramming particle positions from trajectory frames on a fixed 3D grid; here we chose a grid spacing of 1 Å in all directions and all coarse grained particles in the cholesterol molecule were selected for the density calculation. The weighted density for each cluster *k*^′^ was calculated by multiplying the density at each frame belonging to event *t*_*n*_ with the associated probability of the contact event belonging to that cluster, *p*_*n,k*_*′* and subsequent summing of these densities and normalization.

### 2.9 Data Sharing and Software Used

We implemented the Bayesian nonparametric inference from contact time series in the Python package `basicrta` (*Bayesian Single-Cutoff Residence Time Analysis*). Its source code is made available under the open source GNU General Public License, version 3 (or higher) in the GitHub repository github.com/Becksteinlab/basicrta and the latest version is archived under DOI 10.5281/zenodo.13877224. Trajectory data are available through OSF.io at osf.io/aj5wg with DOI 10.17605/OSF.IO/AJ5WG under the Creative Commons Attribution 4.0 International Public License.

In this work we used Python 3.10 with the following packages: MDAnalysis 2.7.0, ^72^ NumPy 1.26,^79^ matplotlib 3.9,^84^ scikit-learn 1.5, ^82^ scipy 1.14,^85^ and tqdm 4.66.^86^ The `basicrta` package used the MDAnalysis MDAKits cookiecutter template version 0.1.^87^ Molecular images were produced with VMD 1.9.3^83^ and Chimera 1.18.^71^

## 3 Results and Discussion

We validated our approach with synthetic data and then applied it to the protein-cholesterol interactions for six different GPCRs.

### 3.1 Validation with synthetic data

To validate our approach, we generated and analyzed a synthetic time series with a set of known parameters (**π** = (0.9, 0.09, 0.01), **λ** = (5, 0.5, 0.05) ns^−1^) that differed from each other by roughly one order of magnitude. The number of exponential components was set to *K* = 15 even though only three were used to generate the 5 × 10^4^ data points. The weights and rates for all components from the Gibbs sampler were clustered with a Gaussian Mixture model as described in Methods. Three well-defined clusters were visible with two diffuse ones superimposed (figure S9 in Supplementary Information). After the latter two clusters were identified as noise (see Methods) and removed, the data in the three remaining clusters provided samples of the posterior distribution, split by component (figure S10 in Supplementary Information). The resulting parameters obtained from the clusters were 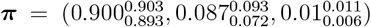, and 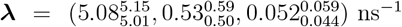, where the reported value is the posterior maximum and the subscript and superscript indicate the lower and upper bounds of the 95% confidence interval. The true values for the second and the important third—the slowest—component were within the confidence intervals while the first, the fastest component, was just outside. Given that in the present application we are primarily interested in the rare slowest component, small deviations in the fastest component are acceptable. As an additional visual check we compared the survival function computed from the synthetic data with the exponential mixture components determined from the clustering results (figure S11 in Supplementary Information) to show how well the parameters determined in the Bayesian analysis fit the input data. In particular, the slowest component matches well the tail of the survival function that contains the few long binding events. In summary, our validation shows that our Bayesian nonparametric approach can infer exponential components that are separated by an order of magnitude given sufficient data set size, and correctly recover the number of components. In the case of the synthetic data with the parameters given above, the number of data points required to correctly distinguish all three components present is on the order of 10^3^.

### 3.2 *β*_2_AR and A_2A_AR

We first applied our Bayesian nonparametric analysis to two GPCRs with high-resolution structures containing resolved cholesterol molecules, namely the beta-2 adrenergic receptor (*β*_2_AR, resolution 2.4 Å)^57^ and the A2A adenosine receptor (A_2A_AR, resolution 1.8 Å).^56^ Although the presence of cholesterol near specific protein residues in crystal structures does not necessarily imply a functional relevance and likely also depends on the detailed experimental conditions, these contacts are nevertheless indicative of potential protein-lipid interactions that may also occur in MD simulation. Here we map high-residence time sites on *β*_2_AR and A_2A_AR and compare our findings to the cholesterol-residue interactions in the crystal structures and to specific interactions that have been highlighted in the literature.

#### *β*_2_AR

We will first show detailed results for a specific residue, W313 in *β*_2_AR, to provide a sense of the numerical results that we will otherwise summarize as a single number, the *τ* MAP estimate from the slowest cluster together with its 95% confidence interval derived from the posterior.

The weights and rates for W313 cleanly separate into five clusters (figure 4) after noise cluster removal (see figure S12 in Supplementary Information) with the slowest timescale of 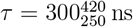; the other clusters span the range from 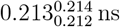 to 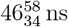 (figure 4). Thus, there exists a clear separation of time scales in the data and the rare, long binding events are clearly distinguishable from faster events. The exponential components with the inferred rates fit the survival function computed from the residence time data fairly well across all observed timescales, from nanoseconds to microseconds (figure 5). The posterior distributions of the weight and rate of the cluster with the slowest rate clearly differ from the priors (figure 6A, B), indicating that the number of data points is sufficiently large so that the shape of the posterior is ultimately determined by the likelihood in Eq. (3) (and not the prior). Thus, given sufficiently large number of contact events (on the order of 10^5^ for our data sets), we can robustly learn the model parameters.

**Figure 5:**
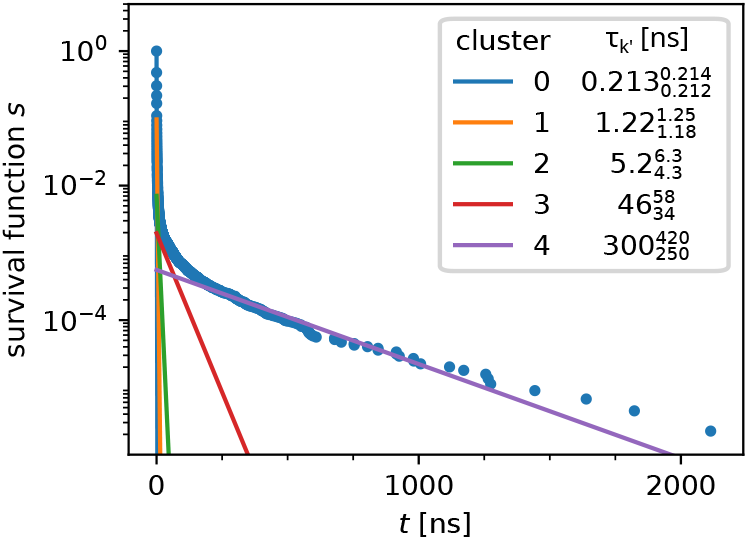
Survival function computed from residence time distribution for the test data set (*β*_2_AR W313) with exponential model components, where exponential mixture model parameters were computed from clusters depicted in figure 4.

**Figure 6:**
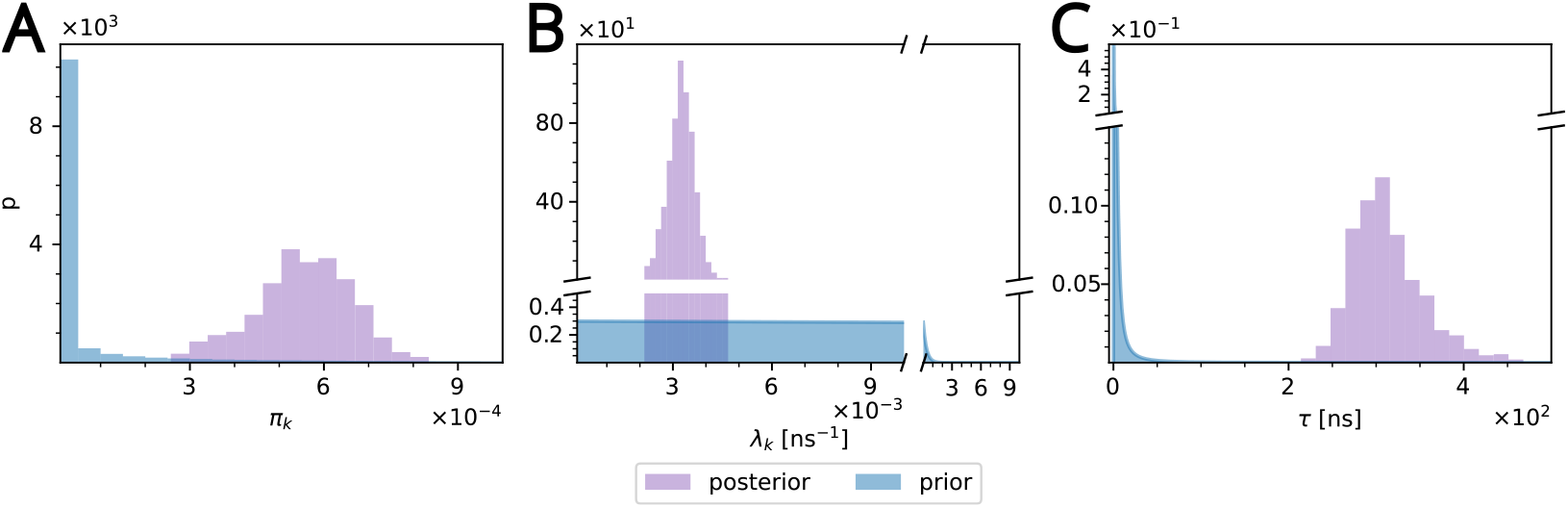
Comparison of posteriors (purple) with priors (blue) for the slowest process in the *β*_2_AR W313 data set (cluster 4 in figure 4). (A) Weight (*π*_*k*_). (B) rate (*λ*_*k*_). (C) The posterior for the residence time (*τ* = *λ*^−1^) was computed by histogramming the transformed rates. The gamma prior (Gamma(*λ*_*k*_; *α*_*k*_, *β*_*k*_)) for the rates becomes an inverse gamma distribution for *τ* (InverseGamma(*τ*; *α*_*k*_, *β*_*k*_)).

In the same manner, the maximum of the slowest waiting time posterior together with its 95% confidence interval was calculated for every single residue. The resulting per-residue *τ* values were projected onto the starting experimental structure to show the regions of long cholesterol interactions in the context of the structure (figure 7A) and plotted as a “fingerprint” for each residue (figure 7B). Although for *β*_2_AR residues with long residence time did not correlate with residues close to cholesterol observed in crystal structures, some of the residues with long residence times have been previously implicated in cholesterol binding: In atomistic MD simulations by Genheden et al. ^88^, the highest-affinity cholesterol binding site for *β*_2_AR was a cleft between I43, G320, and F321, which also includes the long-residence time residue V317. In the same paper, another cholesterol binding site was observed in the cleft between TMs 3, 4, and 5, where V157, V160, and V206 were included in the binding site, all of which are residues with long residence times in our analysis. Another study using atomistic MD observed a cholesterol binding site at the extra-cellular ends of TMs 1 and 7, ^89^ which includes W313 and V317 that were highlighted by our analysis. A cholesterol docking analysis identified P88, C125, V126, F133, I153, V160, T164, V206, W313, and V317 as part of cholesterol binding sites,^90^ all of which were also identified in our coarse grained MD simulations. Another study, which also used coarse-grained MD, observed cholesterol interactions with TM4, which included non-polar interactions with I153 and V157. ^91^ These results indicate that our coarse-grained simulations together with our Bayesian nonparametric analysis of binding event times provides results consistent with previous literature on the interactions of cholesterol with *β*_2_AR.

**Figure 7:**
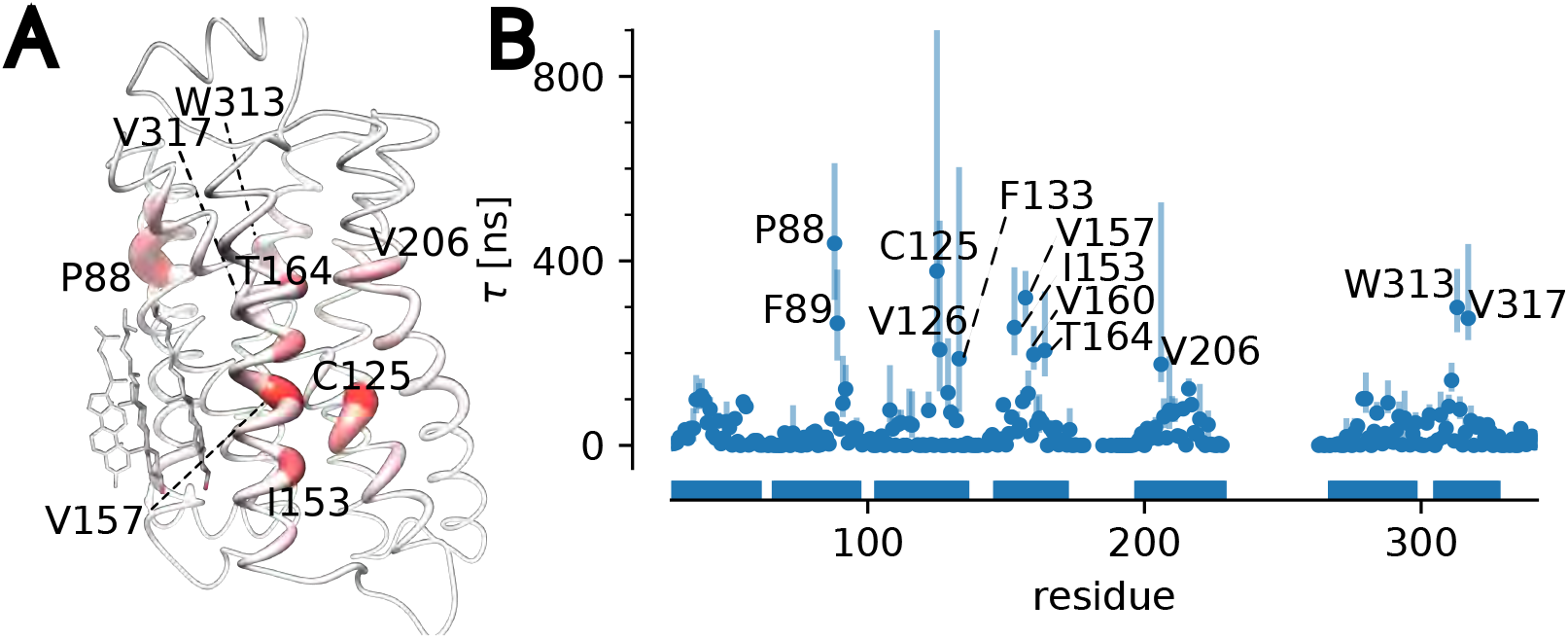
Longest cholesterol residence time *τ* for *β*_2_AR. (A) *τ* projected on the molecular structure. Stronger red color and greater diameter indicate larger *τ*. Cholesterol molecules are resolved in multiple *β*_2_AR structures in generally the same position with PDB 2rh1 shown here serving as as representative structure. (B) *τ* for each residue was determined as the maximum of the posterior distribution (MAP) of the cluster with the smallest rate; error bars indicate the 95% confidence interval. Only residues where *τ* was at least four times larger than the average residence time calculated over all residues are labeled. Horizontal bars indicate the locations of transmembrane helices in the sequence of residues.

#### A_2A_AR

We analyzed cholesterol binding to A_2A_AR in the same manner. The residence times for the slowest process for all residues were projected onto the crystal structure PDB 4eiy and *τ* was plotted against residue for the whole protein. (figure 8). Two residues picked out by the analysis, P248 and L269, match the residues bound to cholesterol in the experimental structure. Other notable residues are F93, A97, and I100, which form a single site to which cholesterol can bind. This site was identified as a region of interaction with coarse grained and all-atom MD simulations.^78^ A docking analysis identified F93, A97, and I100 as part of a cholesterol interaction site, as well as V57, P248, L269, and V275, ^90^ all of which were also highlighted by our analysis. Residues V57, F93, A97, I100, P248^78,92^ and L269^78^ were also shown to interact with cholesterol in MD simulations. Additionally, MD simulations by Genheden et al. ^88^ identified F93, along with other residues, that line the same cleft as A97 and I100. Taken together, our results for A_2A_AR provide a picture consistent with both computational studies and structural data, and predict residues for which cholesterol interactions may be of functional importance.

**Figure 8:**
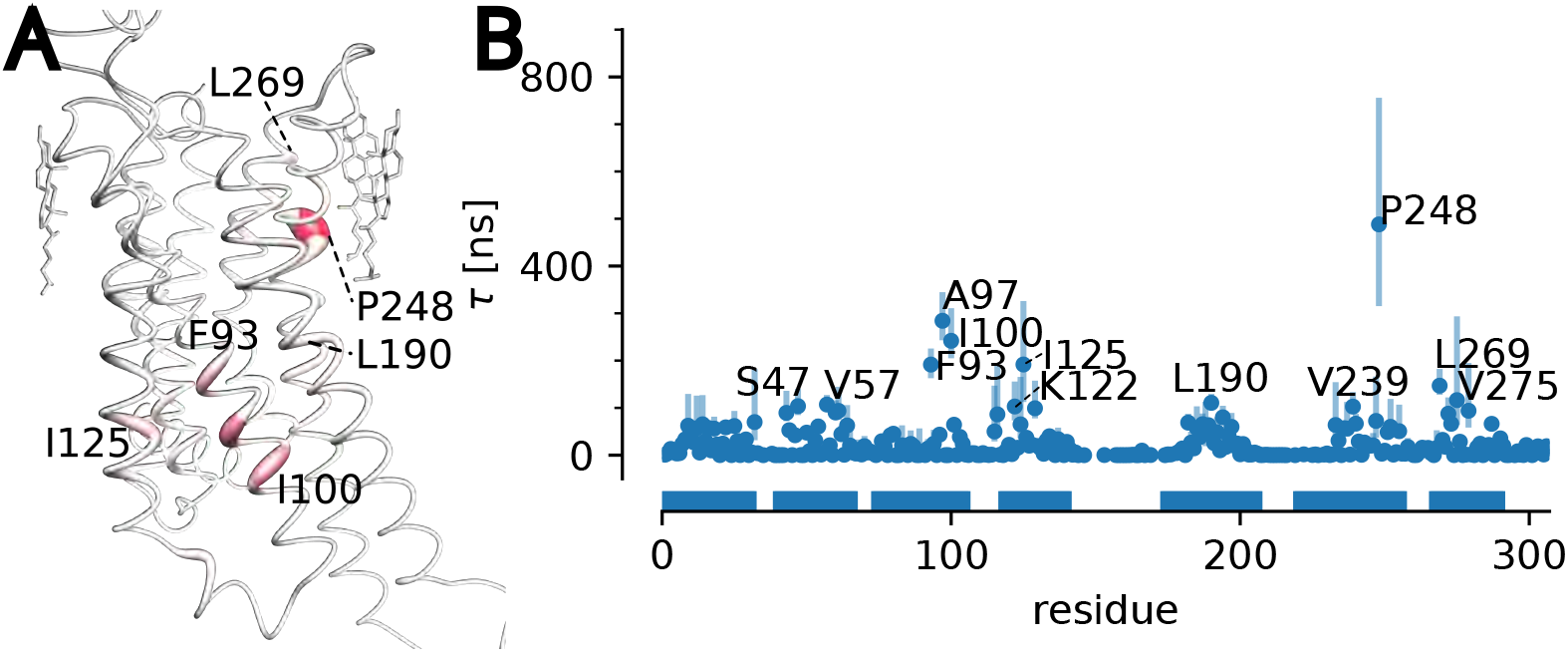
Longest cholesterol residence time *τ* for A_2A_AR. (A) *τ* projected on the molecular structure. Cholesterol molecules shown are resolved in PDB 4eiy. (B) *τ* for each residue (MAP estimate with error bars indicating the 95% confidence interval).

### 3.3 Application to homologous GPCRs

Closely related pairs of GPCRs have been shown to exhibit differences in sensitivity to cholesterol^7–10,30,31^ but it has not always been clear if specific protein-cholesterol interactions are relevant. In the following we investigate the cannabinoid receptors CB_1_R and CB_2_R and the cholecystokinin receptors CCK_1_R and CCK_2_R.

#### Cannabinoid receptors

The cannabinoid receptors CB_1_R and CB_2_R have been shown to respond differently to membrane cholesterol, where CB_1_R activity is diminished with membrane cholesterol enrichment^9^ and CB_2_R is insensitive to membrane cholesterol,^17^ though in one case specific ligands were shown to make CB_2_R cholesterol sensitive.^16^ These two have a sequence similarity of 44% overall and 68% in the transmembrane domain.^96–98^

Per residue *τ* values were determined for both CB_1_R and CB_2_R and projected onto the molecular structures (figure 9). Several residues in contact with cholesterol in the experimental structures of CB_1_R were picked out by the analysis, namely L142, H154, S158, and L286. Residues T201, K232, I243, and V282 were also in contact with cholesterol (although these resides were not labeled in the figure because we only show residues with *τ* of at least four times the mean of all *τ*). No cholesterol were observed in the experimental structures near F381, M384, or L387, which all exhibited large *τ* > 200 ns, but these three resides lined the same cleft, suggesting a cholesterol binding site exists at the intracellular ends of TMs 1 and 7. In CB_2_R, residues L126, P260, F283, C284, and M286 were all found near experimentally resolved cholesterol. L71, S75, and V86 (high *τ* > 150 ns) in TM2 and K109 in TM3 face the same cleft, indicating a region where cholesterol interactions may be important.

**Figure 9:**
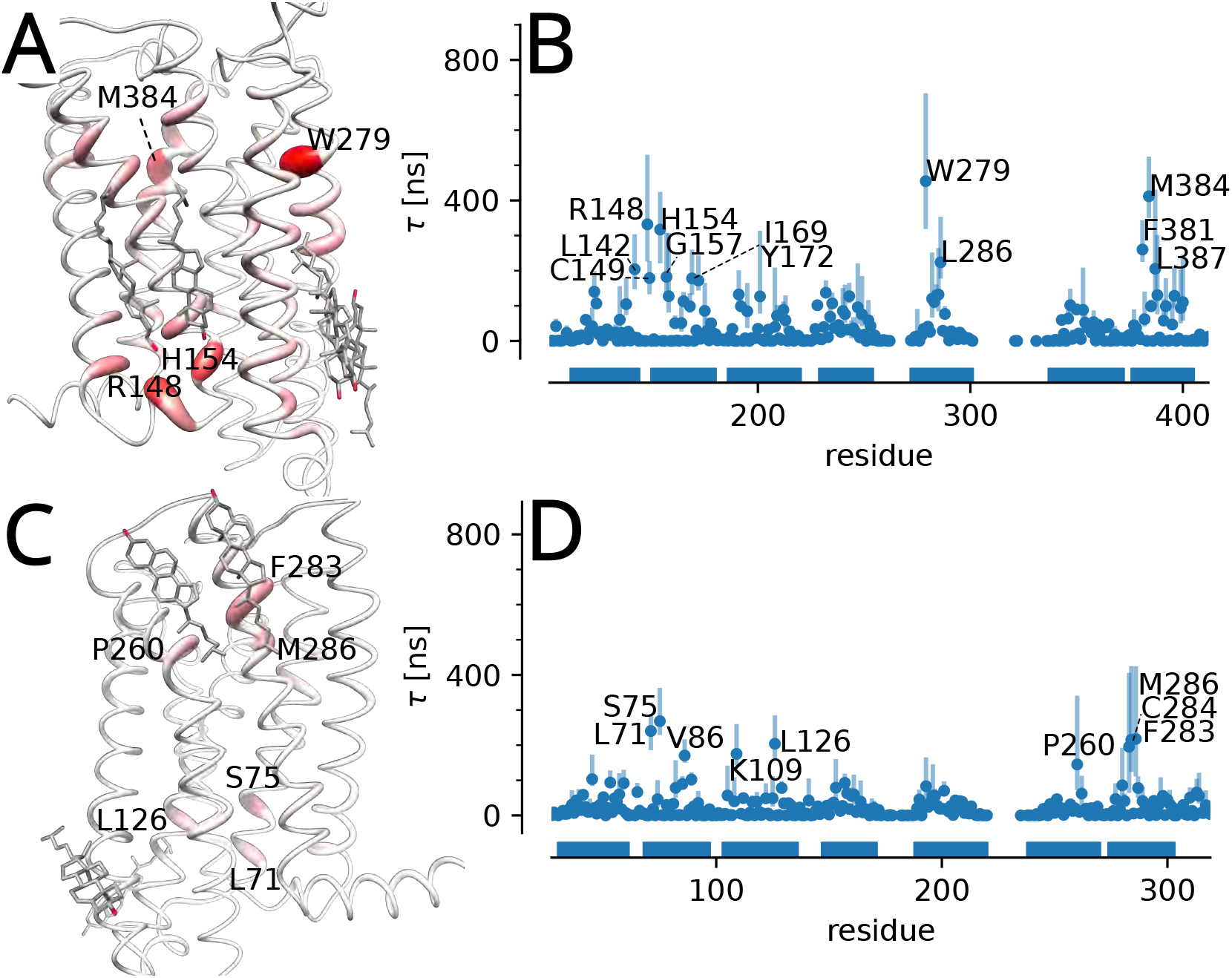
Longest cholesterol residence time *τ* for the cannabinoid receptors. (A) *τ* for CB_1_R projected on the molecular structure. Cholesterol molecules from the structures PDB 5xr8^93^ and PDB 6n4b^94^ are shown. (B) *τ* for each residue in CB_1_R (MAP estimate with error bars indicating the 95% confidence interval). (C) *τ* projected on CB_2_R (PDB 6pt0^95^). (D) *τ* for each residue in CB_2_R.

In order to compare *τ* between equivalent sites on the proteins, 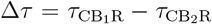 was computed for each matching pair of residues in the sequence alignment. Δ*τ* is generally large and positive, indicating longer interaction times for CB_1_R than for CB_2_R at equivalent sites, especially W279, R148, and M384 (figure S14A in Supplementary Information). The same conclusion holds when the interaction of cholesterol with any part of the protein is analyzed, as shown by the whole-protein survival function, which was computed from the combined set of contact times from all residues for each protein (figure S16A in Supplementary Information). The survival function decays faster for CB_2_R than for CB_1_R, indicating that the probability to observe long interaction times is smaller for CB_2_R than CB_1_R. The results for CB_1_R and CB_2_R show that the Bayesian nonparametric analysis of binding event times is able to highlight specific cholesterol-residue interactions that are consistent with experimental structures. The global analysis of all residues indicates that the more cholesterol-sensitive homolog, CB_1_R, contains more residues with longer residence times than CB_2_R, the cholesterol insensitive one.

#### Cholecystokinin receptors

The cholecystokinin receptors CCK_1_R and CCK_2_R have a sequence similarity of 53% overall and 69% in the transmembrane domain.^7^ Although both receptors have cholesterol recognition amino acid consensus (CRAC) motifs in TMs 3 and 5 and a cholesterol consensus motif (CCM) in TM 4, these proteins exhibit different levels of sensitivity to membrane cholesterol.^7^ Signaling and ligand binding in CCK_1_R both decrease upon cholesterol depletion and enrichment,^30^ and the signaling of CCK_2_R is affected little, if at all, by membrane cholesterol.^6,7^ We had previously investigated CCK_1_R and CCK_2_R and found that CCK_1_R contained more specific residues with longer residence times than CCK_2_R;^50^ here we are revisiting the cholecystokinin receptors with additional simulations and our new Bayesian nonparametric approach to provide an updated evaluation of the source of cholesterol sensitivity in these two homologous GPCRs.

Per residue *τ* values were determined for both CCK_1_R and CCK_2_R (figure 10) and projected onto the models. Currently no structures of CCK receptors include resolved cholesterol, but one high-resolution electron microscopy structure of CCK_1_R (PDB 7mbx) includes a resolved cholesterol hemisuccinate molecule. Two residues with high *τ* values in CCK_1_R, I67 and I78, are near the resolved cholesterol hemisuccinate in 7mbx. The remainder of the residues labeled in figure 10B (with the exception of I368) all line the same cleft between TMs 3, 4, and 5, indicating a cholesterol binding pocket. In both proteins, five equivalent residues form the majority of the binding site (S126, F130, V133, A134, L137 in CCK_1_R and S139, L143, V146, A147, L150 in CCK_2_R), while the other residues lining the same cleft are identical for both proteins (L168, S169 in CCK_1_R are equivalent to L181, S182 in CCK_2_R and I229, P230 in CCK_2_R are equivalent to I220, P221 in CCK_1_R). A previous study of cholesterol sensitivity of these two homologs^7^ revealed a mutation which renders CCK_1_R insensitive to cholesterol, namely Y140A, which lines the same TM 3-4-5 binding pocket and is one helical turn from L137. These results may indicate that this binding pocket in CCK_1_R is important for cholesterol sensitivity. Our analysis also revealed a second cholesterol binding site in CCK_2_R between TMs 2 and 4, which includes S95, S99, I175, V175, and W179. This binding site is absent in CCK_1_R, which may indicate that it is not relevant for the differential cholesterol sensitivity between CCK_1_R and CCK_2_R.

**Figure 10:**
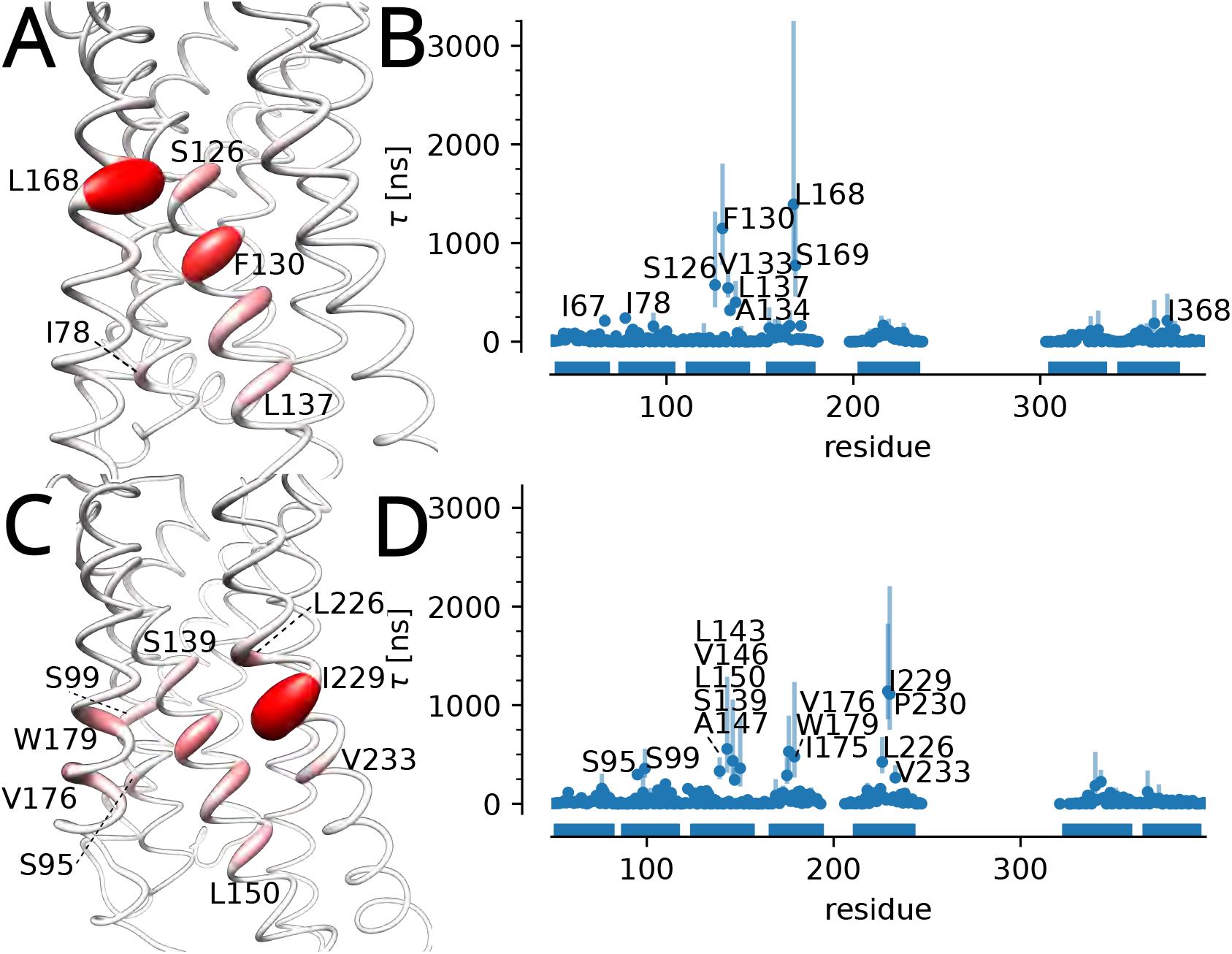
Longest cholesterol residence time *τ* for the cholecystokinin receptors. (A) *τ* for CCK_1_R projected on the molecular model of CCK_1_R. (B) *τ* for each residue in CCK_1_R (MAP estimate with error bars indicating the 95% confidence interval). (C) *τ* projected on the model of CCK_2_R. (D) *τ* for each residue in CCK_2_R. Residues 235 to 313 for CCK_1_R and 243 to 333 in CCK_2_R form a cytoplasmic domain that was not included in the models.

As for the cannabinoid receptors, we computed the difference 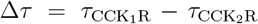 for each matching pair of residues in the sequence alignment (figure S14B in Supplementary Information). Δ*τ* is large and greater than zero for residues in CCK_1_R (especially L168, F130, and S169, which we also highlighted in our previous work^50^). The major difference compared to Geiger et al. ^50^ is the detection of a second binding site formed by V176, W179, and S99 in CCK_2_R, which is apparently absent in CCK_1_R. Overall, CCK_1_R displays similar interaction residues and *τ* when compared to those of CCK_2_R, which is corroborated by the similar whole-protein survival functions (figure S16B in Supplementary Information). These results suggest that CCK_1_R may be more sensitive to cholesterol due to subtle differences in the cholesterol-binding site interactions and not simply due to overall longer cholesterol binding. The larger *τ* values at the TM 3-4-5 binding site in both CCK_1_R and CCK_2_R indicate this region may be important for cholesterol sensitivity, but our simulations do not reveal a mechanism of cholesterol modulated function. A more detailed analysis of the specific cholesterol-protein interactions at the indicated interaction sites may be useful in further understanding the role cholesterol plays in modulating protein function, and a first step in this direction is our *kinetic mapping* approach that allows us to associate residence times with binding poses.

### 3.4 Kinetically mapped cholesterol binding modes

Our Bayesian nonparametric approach allows us to associate frames in the underlying MD trajectory with clusters and thus time scales of binding events. One of our underlying motivating assumptions for the exponential mixture model Eq. 1 was that different exponential components would correspond to different binding processes. In the following we show examples of analyzing the MD trajectories “by component” and see that indeed different components (and thus time scales) describe physically distinct processes. We focus on the quantitative description of localization of cholesterol in the binding site by the weighted 3D density and the qualitative visualization of the binding modes by molecular graphics.

#### Weighted density

We can use the probability of a trajectory frame to have contributed to a cluster (see Eq. 10 in Methods) to compute a cluster-weighted density of cholesterol over the trajectory. For binding of cholesterol to W313 in *β*_2_AR (see figure 4), the weighted density (computed over all trajectory frames in which cholesterol binding was observed) shows distinct localization of cholesterol relative to the W313 residue, depending on the associated residence time *τ*_*k*_*′* (figure 11). For the two fastest rates (corresponding to the shortest residence times, 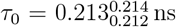 and 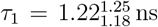) the densities are more diffuse and located farther away from W313 than for the binding events associated with slower rates, which becomes especially clear when the isocontour cutoff for the density depiction is increased. The densities for the two components with the slowest rates (longest residence times, 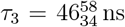 and 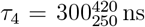) were more localized near the residue W313. The orientation of the W313 sidechain is the same for the slow processes 3 and 4 and differs from the faster components 0, 1, and 2. Thus, the kinetic mapping not only picked up different cholesterol localizations but also distinct changes in the protein binding site that were associated with the stable binding mode.

**Figure 11:**
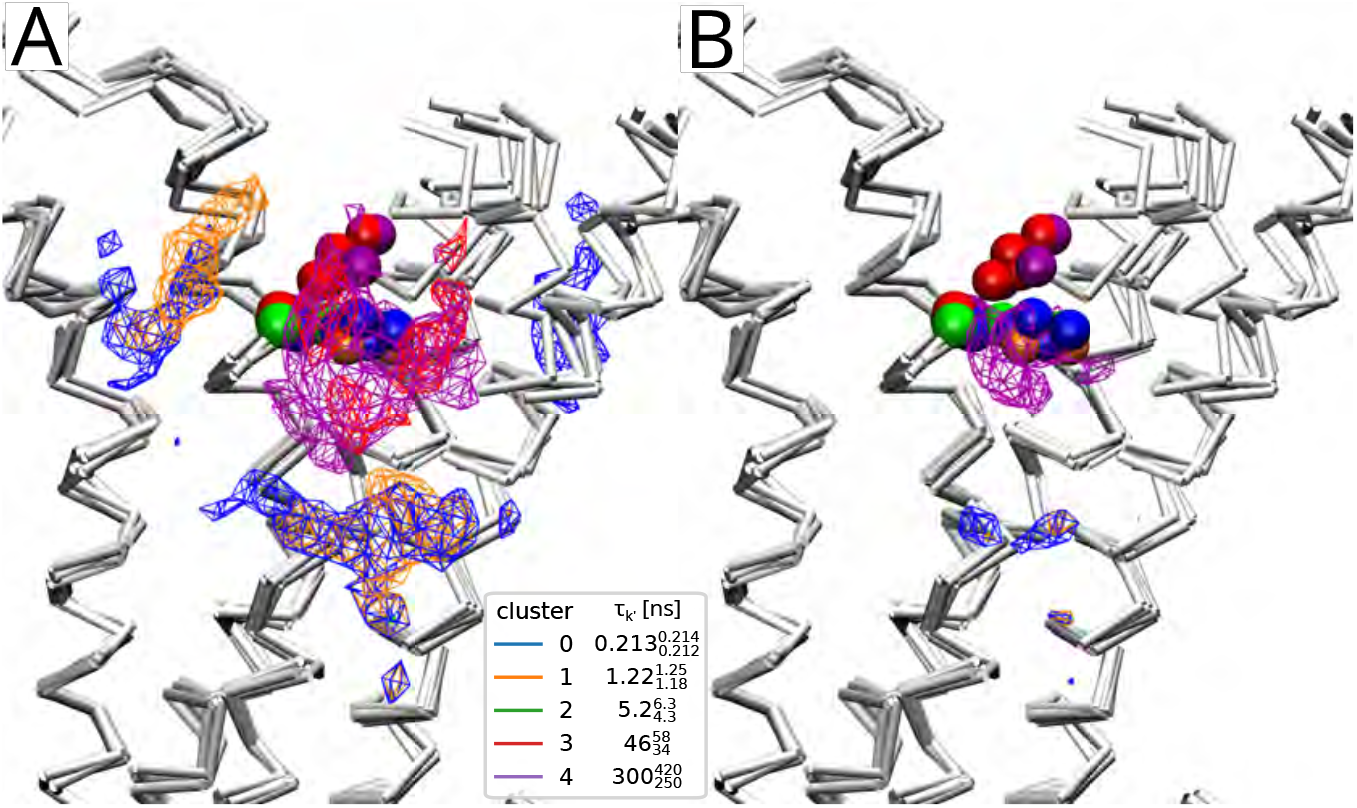
Weighted densities for W313 in *β*_2_AR, where W313 is shown in the VDW representation and the colors correspond to the result plot in figure 5. (A) Densities shown at 0.002 Å^−3^ isocontour cutoff show regions of density near the cutoff corresponding to the fastest rate (blue) and second fastest rate (orange). The second slowest (red) and slowest (purple) components are more localized near the W313 but have different shapes due to the differences in binding events. The density for cluster 2 (green) is not visible at this isocontour cutoff. (B) Densities depicted at 0.005 Å^−1^ show the most dense region near W313 is due to the slowest process, where the sidechain rearranges and allows for cholesterol to assume a conformation resulting in longer bound times.

The same analysis was performed for two other residues in *β*_2_AR (P88, see figure S17 for the clustering, and V157, see figure S18 in the Supporting Information). The weighted density for P88 (figure S19 in Supplementary Information) also shows that for the component with the largest rates 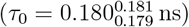, cholesterol is localized away from P88 near the range of the cutoff (7 Å). The three components with the slowest rates are localized near the residue 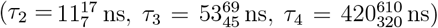, with the density of the slowest component (*τ*_4_) slightly closer to the protein. Component 4 differs from the other components in that the sidechain of F89 changed conformation and makes contact with cholesterol (figure S19 in Supplementary Information), which may be partially responsible for the longer lasting binding events. The weighted density for V157 (figure S20 in Supplementary Information) show the same pattern for densities of the fast components 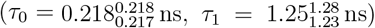 as they are located at the periphery of V157. The density of the slowest component 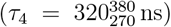 is located closest to V157 and occupies a well-defined elongated region inside a cleft between helices 3, 4, and 5.

The weighted densities quantify the cholesterol positions in the kinetically mapped trajectories and show that cholesterol localization depends on the time scale of the binding events. Additionally, distinct sidechain conformations are seen to be associated with the long timescale binding events. Although only weighted densities were analyzed in this work, in principle any number of analyses could be performed on the kinetically mapped trajectories, such as geometric clustering, interactions with solvent, or detailed ligand-protein interactions.

#### Binding poses

The weighted density analysis indicated that cholesterol would likely assume different binding poses depending on the time scale of the binding process. We use molecular graphics with VMD^83^ with the kinetically mapped trajectory to directly visualize such differing poses.

Residue M384 in CB_1_R had been identified as one of the ones with the longest cholesterol residence time (figure 9B and figure S13A in Supplementary Information) so we wanted to visualize the difference between the slowest process 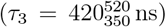 and the second slowest one 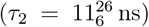. Five hundred frames with the greatest probability of belonging to each process were drawn on top of each other in VMD^83^ to visualize the binding pose (figure 12). The second slowest process (cluster 2) shows greater variability in the position of the polar end of cholesterol (figure 12B), which may indicate several binding modes with similar off-rates. The slowest process (cluster 3) displays a much more consistent binding mode across frames, where the polar end of cholesterol is inserted in a pocket formed near S123, F381, and M384 (figure 12C). The visualization demonstrates that different components in the exponential mixture model (Eq. 1) correspond to different binding modes and thus, these different modes lead to binding time scales (and off rates) that differ by more than an order of magnitude.

**Figure 12:**
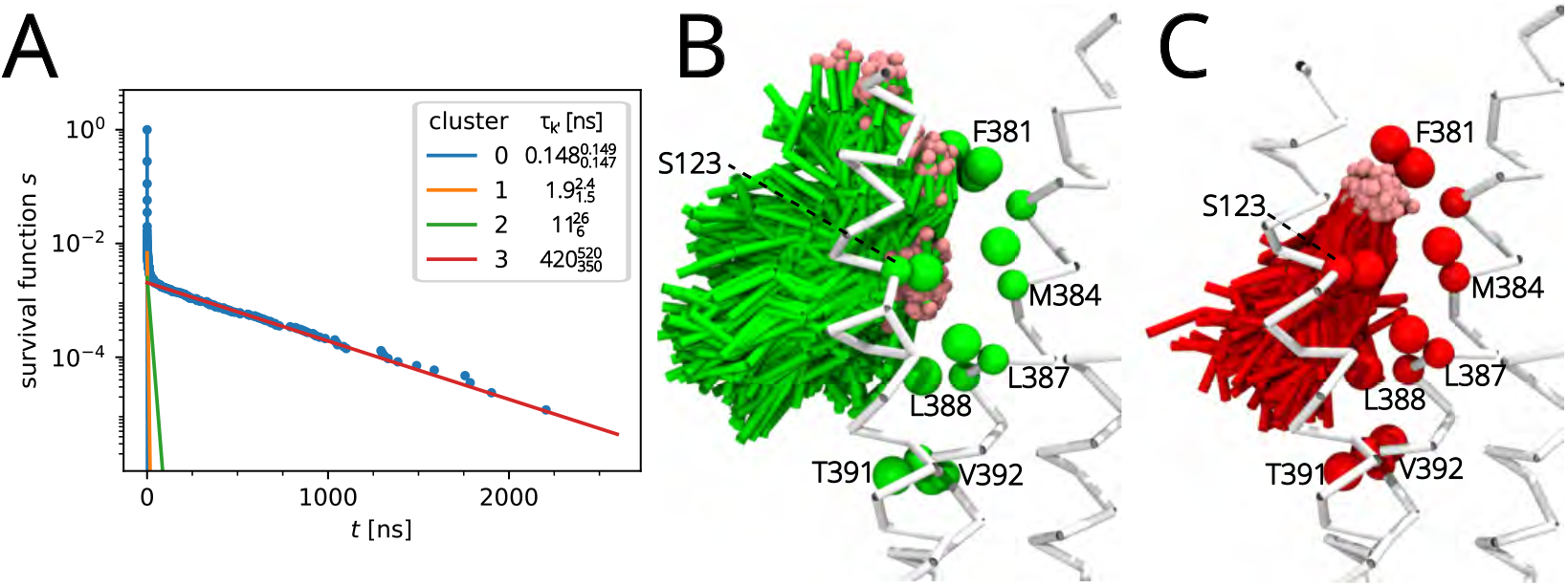
Cholesterol binding to M384 of CB_1_R. (A) Survival function of the data (blue dots) with individual cluster components *k*^′^ (0–3) with inferred residence times *τ*_*k*_*′* (plotted as individual exponential components). Detailed clustering of samples is shown in figure S13A in Supplementary Information. (B) Binding poses of cholesterol associated with the second slowest cluster *k*^′^ = 2 (green) with 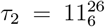 as obtained with the kinetic mapping approach. (C) Binding poses associated with the slowest cluster *k*^′^ = 3 (red) with 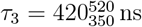. The pink sphere indicates the polar group in the coarse-grained cholesterol model.

Residue M384 in CB_1_R is equivalent to M286 in the homologous receptor CB_2_R. We therefore compared the binding mode of M286 to that of M384 to understand if equivalent positions lead to similar binding modes. As figure S21B,D in Supplementary Information show, cholesterol adopts a well-defined binding pose for each of the slowest processes for M384 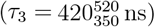 and M286 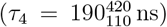 but the location of the cholesterol hydroxyl group and the overall orientation differs substantially. In CB_1_R, cholesterol is approximately parallel to the membrane normal with the hydroxyl located near polar S123. In CB_2_R, its orientation is nearly in the plane of the membrane and the hydroxyl is close to M286. Evidently, the contacts formed in CB_1_R result in longer binding events although a more detailed analysis is needed to determine precisely the cause of the increased affinity for cholesterol near M384. This initial analysis shows that equivalent residues in homologous proteins do not necessarily lead to similar binding modes of cholesterol.

A similar analysis was performed for the residues that form the TM 3-4-5 binding site in CCK_1_R and CCK_2_R, which showed different binding poses for the longest binding events in each protein (figure S15 in Supplementary Information). In CCK_1_R, cholesterol binds between TM 3 and TM 4 but in CCK_2_R it binds between TM 3 and TM5. F130 in CCK_1_R appears to clash with the cholesterol in CCK_2_R, thus preventing a binding pose similar to that of CCK_2_R. Experimentally, the Y140A mutation abolishes cholesterol sensitivity in CCK_1_R^7^ and given the location of Y140 (in CCK_1_R) at the bottom of the equivalent CCK_2_R binding site, it is possible that the mutation removes a steric clash at the tail of the cholesterol that may allow for a more CCK_2_R-like binding pose in CCK_1_R and thus remove the interaction with TM 4. Although a more detailed analysis of the interactions is needed, the present analysis hints at F130 in CCK_1_R as important for the differential cholesterol sensitivity between the two homologs.

## 4 Conclusions

We introduced a Bayesian nonparametric analysis for contact time series to determine the residence time *τ* for an interaction of interest. As the underlying model we chose a *K*-term exponential mixture model (Eq. (1)), based on the hypothesis that an unknown number of different, simple molecular processes, each represented by a single exponential component, contribute to contacts at a given site. The introduction of a latent variable that maps samples to components led to simple expressions for the posterior distribution of the weights and rates of the exponential components. Importantly, in contrast to a model-selection scheme based in the Akaike or Bayesian Information Criteria (AIC or BIC respectively) that is only asymptotically valid in the large-data limit, the Dirichlet process prior we place on weights^51^ allows us to rigorously learn the number of components alongside their rates regardless of the data set size. The posterior was numerically sampled with a Markov Chain Monte Carlo Gibbs sampler while the latent variable allowed for the probabilistic mapping of the components with their associated kinetic rates back to the samples (*kinetic mapping*). When the contact time samples were obtained from a MD trajectory, we could assign frames of the trajectory to specific components and in this way cluster the trajectory by the timescales (set by the inverse rates of the components) of the binding process of interest. In this way it is possible to map back the coarse observable (the contacts) to the underlying molecular processes.

The Bayesian approach was first validated with synthetic data, where the number of components and their values, and in particular the “rare” component with low probability and small rate, were recovered. Motivated by our earlier work,^50^ we returned to the problem of finding residues on GPCRs that interact preferentially with cholesterol in the membrane. Such residues were detected from the cholesterol-residue contact time series as those with the smallest off-rate (or equivalently, longest residence time) in the Bayesian analysis. In total we analyzed over 1 ms of coarse-grained MD trajectories for six different GPCRs. In three of four GPCRs with experimentally resolved cholesterol-protein crystal structure contacts, the simulations predicted ten structurally confirmed specific protein-cholesterol interactions (for A_2A_AR, CB_1_R, CB_2_R); for *β*_2_AR (and A_2A_AR), our results are consistent with other computational studies. In the homologous pair of receptors CB_1_R and CB_2_R, more residues with longer residence were predicted for CB_1_R, which may be part of the explanation why CB_1_R has been experimentally found to be sensitive to membrane cholesterol while CB_2_R has been insensitive. Simulations for a similarly related pair, CCK_1_R and CCK_2_R, showed little difference in overall binding even though CCK_1_R is known to be more cholesterol sensitive than CCK_2_R. However, a more detailed analysis of the cholesterol binding poses from the kinetic mapping showed differences in cholesterol binding sites due to differences in specific residues and suggested a potential mechanism for the Y140A mutation that abolishes cholesterol sensitivity in Y140A CCK_1_R. Although more work is required, these initial results indicate that in the case of the cholecystokinin receptors, cholesterol sensitivity may be due to very specific interactions.

In order to test our underlying hypothesis that different components in our model represent different distinct molecular processes, we used the kinetic mapping approach to analyze binding to selected residues by their associated timescale. Visualization with frames clustered by the most probable component showed distinct cholesterol binding poses and in some cases, different conformers of the residue in question. The weighted 3D density of cholesterol around the target residue confirmed the visual analysis quantitatively and showed that binding modes with long timescales tend to be closer to the residue and the protein surface and often have distinct shapes compared to the faster (short time scales) binding modes. These fast modes tend to have more diffuse densities on the edge of the cutoff or overlap with binding modes associated with neighboring residues, thus indicating that they may be due to fluctuations around the cutoff distance. Thus, our Bayesian nonparametric analysis of binding with an exponential mixture model for the residence time distribution is able to filter out the functionally irrelevant or artificial short binding events while quantifying the relevant long binding events.

Given the use of MD simulations in the elucidation of important interactions between proteins and their local environments, we provide a robust method for the analysis of MD contact time series to infer the longest residence times for the rare binding events with quantitative determination of binding sites on a protein. Such time series could be obtained with more sophisticated algorithms than the single-cutoff thresholding that we used; for example, a double cutoff comparator with hysteresis (a Schmitt trigger^99,100^) to reduce the number of fluctuations across the cutoff or a time filter that ignores intermittent loss of contact for a short duration^101^ have been used in other contexts. We focused on GPCR-cholesterol interactions, but our approach is not limited to protein-lipid interactions and can in principle be used to analyze any protein-small molecule interactions from MD simulations where a sufficient number of binding/unbinding events can be observed. As such, it may find broader applications in computer-aided drug discovery and molecular pharmacology. The Bayesian non-parametric approach yields full posteriors and thus enables a rigorous assessment of errors. The kinetic mapping approach that is inherent in our construction of the posteriors provides a new approach to associate timescales of processes that were measured by a macroscopic observable such as a simple distance to the underlying microscopic processes and mechanisms. Our method may thus aid in inferring accurate kinetics from any contact-like time series and—if the time series was sampled with MD simulations—discovering molecular mechanisms associated with binding of ions, lipids, and small or drug-like molecules to proteins and nucleic acids.

## Acknowledgement

Research reported in this publication was supported by the National Institute of General Medical Sciences of the National Institutes of Health under Award Number 1R15GM124623 (O.B.). S.P. acknowledges support of the NIH (Grant No. R35GM148237). The authors also acknowledge Research Computing at Arizona State University for providing HPC and storage resources through Agave, Phoenix, and Sol, ^102^ which have contributed to the research results reported within this paper.

## Supporting Information Available

The Supporting Information is available free of charge at https://.

