## Supplementary Information for "Bayesian nonparametric analysis of residence times for protein-lipid interactions in Molecular Dynamics simulations"

**Abstract**

The Supplementary Information contains additional details for the main paper. We include an example of using least squares fitting, the current state-of-the-art for protein-lipid interaction analysis, for the $\beta_2$AR W313 data set to demonstrate the shortcomings of this method compared to our new Bayesian nonparametric inference approach. We include details for the derivation of the conditional posterior probabilities that are used for the Gibbs sampler algorithm. We analyzed the sensitivity of our approach to changes in the cutoff for the contact analysis by varying the cutoff value between 6 Å and 8 Å. The maximum residence times $\tau$ increase with cutoff (especially from 6 Å to 7 Å) while qualitatively leading to the same conclusions by identifying the same long-binding residues. We show that our results are insensitive to the exact choices for burn-in and thinning (subsampling) for processing the samples from the Gibbs sampler. We show additional results for the Bayesian nonparametric analysis of cholesterol binding to GPCRs.

#### S1 Least squares fitting

A common practice in determining model parameters that match the observed data is by minimizing the sum of the squares of the differences between model and data, also known as *least squares fitting*.[1] This approach has been used to estimate waiting times for lipid-protein binding from MD simulations by extracting the waiting time distribution $S(t)$ from the simulations and fitting to an exponential mixture model with fixed number of components $K$[2,3]

$$y(t|\boldsymbol{\pi}, \boldsymbol{\lambda}) = 1 - \int_0^t p(t')dt' = \sum_{k=1}^{K} \pi_k e^{-\lambda_k t}. \tag{S1}$$

Minimizing the sum of the squared residuals with respect to the model parameters $\boldsymbol{\theta}$ (weights $\pi_k$ and rates $\lambda_k$)

$$\hat{\boldsymbol{\theta}} = \arg\min_{\boldsymbol{\theta}} \sum_n \left(S(t) - y(t|\boldsymbol{\theta})\right)^2 \tag{S2}$$

yields point estimates $\hat{\boldsymbol{\theta}}$ of the parameters. Note that the least squares fitting procedure requires a function of the input data ($S(t)$) and does not directly work with the raw data (the waiting times $t$) themselves. This straightforward approach appears to be the current "state-of-the-art" for lipid-protein interaction analysis.[2,3]

In our previous work[2] we used the least squares fitting approach with careful choices of weights to make it slightly more robust, but to the best of our knowledge, this is not how least squares has been commonly used, and hence in the following we show how simple least squares fitting (with equal weights for all data points) compares to our Bayesian nonparametric approach from the main paper. Least squares fits were performed using the *curve_fit*() function of scipy.[4] We focus on our representative data set ($\beta_2$AR W313) as an example.

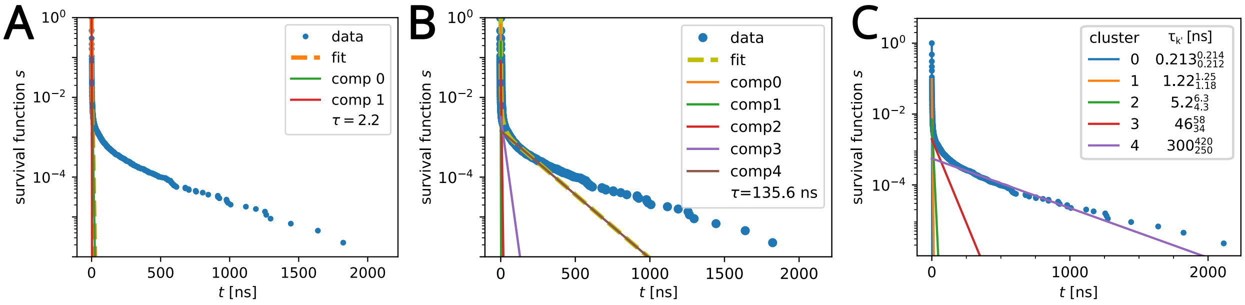


**Figure S1:** Survival function of the representative data set ($\beta_2$AR W313) with 2 component exponential fit using least squares (A), a 5 component fit using least squares (B), and the 15-component nonparametric Bayesian results of the current method (C).

As described in the main paper, our best estimate for this dataset is a $K = 5$ component model with the slowest component $\tau = 300^{420}_{250}$ ns. For the least square fit with $K = 2$, the fit is completely dominated by the overwhelming number of short events as can be seen in the comparison of the survival function of the data to the fit and the individual components (figure S1A). Even if we choose the correct number of components $K = 5$ for the fit, $\tau = 135.6$ ns is still outside the 95% confidence interval of the Bayesian estimate and the fit clearly fails to describe the sample distribution (figure S1B). In our experience, the solutions for this non-linear fitting problem can also be sensitive to the initial conditions of the solver and sometimes do not yield an ideal curve fit. The least squares solution only provides a point estimate of the optimal parameters; error estimates need to be obtained by other methods such as bootstrapping or dividing the trajectory into blocks and performing the analysis on each block.[2] The latter method has the distinct disadvantage that the artificial blocking destroys long events and thus further reduces the amount of rare events sampled while also artificially increasing $\tau$ estimates; choosing the number of blocks that has the lowest effect on the error estimates is difficult.[2]

Our Bayesian method, on the other hand, allows for an arbitrary number of components that best fit the data (figure S1C). It sidesteps the model selection problem of having to choose $K$ in advance and instead provides the components that best describe the data. It provides full joint posterior distributions of the model parameters and thus yields rigorous error estimates that faithfully quantify the uncertainty. Although the generation of the posteriors is computationally more expensive than least square fitting, we found the approach to be robust without the need of human intervention or tuning of parameters, so that it can be easily automated and run for large data sets.

### S2 Conditional posteriors

The likelihood for the model Eq. (1) before the inclusion of the latent indicator is

$$p\left(\boldsymbol{t}|\boldsymbol{\pi},\boldsymbol{\lambda}\right)=\prod_{n=1}^{\Omega}\sum_{k=1}^{K}\pi_k\lambda_k e^{-\lambda_k t_n}. \tag{S3}$$

The introduction of the indicator allows for the classification of each of the $\Omega$ data points with one of the unbinding processes with exponentially distributed residence times. The completed likelihood[5] then becomes

$$p\left(\boldsymbol{t}|\boldsymbol{\lambda},\boldsymbol{z}\right)=\prod_{n=1}^{\Omega}\lambda_{z_n} e^{-\lambda_{z_n} t_n}. \tag{S4}$$

The conditional posteriors for each of the parameters are determined from Bayes' theorem.

##### Indicator

For the conditional posterior of the indicator components $\mathbf{z}=(z_1,\ldots,z_n,\ldots,z_\Omega)$, Bayes' theorem is

$$p\left(z_n|t_n,\boldsymbol{\lambda},\boldsymbol{\pi}\right)p\left(t_n|\boldsymbol{\lambda},\boldsymbol{\pi}\right)=p\left(t_n|z_n,\boldsymbol{\lambda},\boldsymbol{\pi}\right)p\left(z_n|\boldsymbol{\lambda},\boldsymbol{\pi}\right),$$

for which we can write

$$\begin{aligned}p(z_n|t_n,\boldsymbol{\lambda},\boldsymbol{\pi}) &\propto p(t_n|z_n,\boldsymbol{\lambda},\boldsymbol{\pi})p(z_n|\boldsymbol{\lambda},\boldsymbol{\pi})\\ &= p\left(t_n|z_n,\boldsymbol{\lambda},\boldsymbol{\pi}\right)p\left(\boldsymbol{\lambda}|z_n,\boldsymbol{\pi}\right)p\left(z_n|\boldsymbol{\pi}\right)p\left(\boldsymbol{\pi}\right).\end{aligned}$$

Since $\boldsymbol{\lambda}$ is statistically independent of $\mathbf{z}$ and $\boldsymbol{\pi}$, this can be written as

$$\begin{aligned}p(z_n|t_n,\boldsymbol{\lambda},\boldsymbol{\pi}) &\propto p\left(t_n|z_n,\boldsymbol{\lambda},\boldsymbol{\pi}\right)p\left(\boldsymbol{\lambda}\right)p\left(z_n|\boldsymbol{\pi}\right)p\left(\boldsymbol{\pi}\right)\\ &\propto p\left(t_n|z_n,\boldsymbol{\lambda},\boldsymbol{\pi}\right)p\left(z_n|\boldsymbol{\pi}\right)\end{aligned}$$

where $p(t_n|z_n, \boldsymbol{\lambda}, \boldsymbol{\pi})$ is the term in the completed likelihood belonging to data point $n$ and $p(z_n|\boldsymbol{\pi})$ the categorical prior on $\mathbf{z}$. Inserting the likelihood and prior yields

$$p\left(z_n|t_n, \boldsymbol{\lambda}, \boldsymbol{\pi}\right) \propto \pi_{z_n} \lambda_{z_n} e^{-\lambda_{z_n} t_n},$$

which provides a weighted probability for each data point $t_n$ to belong to each of the $K$ components. After normalizing, the conditional posterior of the indicator becomes

$$p\left(z_n|t_n, \boldsymbol{\lambda}, \boldsymbol{\pi}\right) = \frac{\pi_{z_n} \lambda_{z_n} e^{-\lambda_{z_n} t_n}}{\sum_{k=1}^{K} \pi_k \lambda_k e^{-\lambda_k t_n}}. \tag{S5}$$

**Rates**

For the rates ($\boldsymbol{\lambda}$), Bayes' theorem is

$$p\left(\boldsymbol{\lambda}|\boldsymbol{t}, \boldsymbol{\pi}, \mathbf{z}\right) p\left(\mathbf{z}|\boldsymbol{\pi}, \boldsymbol{t}\right) p\left(\boldsymbol{\pi}|\boldsymbol{t}\right) p\left(\boldsymbol{t}\right) = p\left(\boldsymbol{t}|\boldsymbol{\lambda}, \boldsymbol{\pi}, \mathbf{z}\right) p\left(\mathbf{z}|\boldsymbol{\pi}, \boldsymbol{\lambda}\right) p\left(\boldsymbol{\pi}|\boldsymbol{\lambda}\right) p\left(\boldsymbol{\lambda}\right).$$

Since the indicator is statistically independent of both $\boldsymbol{t}$ and $\boldsymbol{\lambda}$ and the weights are statistically independent of both $\boldsymbol{t}$ and $\boldsymbol{\lambda}$, this can be written as

$$p\left(\boldsymbol{\lambda}|\boldsymbol{t}, \boldsymbol{\pi}, \mathbf{z}\right) p\left(\mathbf{z}|\boldsymbol{\pi}\right) p\left(\boldsymbol{\pi}\right) p\left(\boldsymbol{t}\right) = p\left(\boldsymbol{t}|\boldsymbol{\lambda}, \boldsymbol{\pi}, \mathbf{z}\right) p\left(\mathbf{z}|\boldsymbol{\pi}\right) p\left(\boldsymbol{\pi}\right) p\left(\boldsymbol{\lambda}\right).$$

The conditional posterior for the rates $\boldsymbol{\lambda}$ is then

$$p\left(\boldsymbol{\lambda}|\boldsymbol{t}, \boldsymbol{\pi}, \mathbf{z}\right) = \frac{p\left(\boldsymbol{t}|\boldsymbol{\lambda}, \boldsymbol{\pi}, \mathbf{z}\right) p\left(\boldsymbol{\lambda}\right)}{p\left(\boldsymbol{t}\right)}.$$

Since the denominator is independent of $\boldsymbol{\lambda}$ and the completed likelihood Eq. (S4) is independent of $\boldsymbol{\pi}$, it can be written as

$$p\left(\boldsymbol{\lambda}|\boldsymbol{t}, \mathbf{z}\right) \propto p\left(\boldsymbol{t}|\boldsymbol{\lambda}, \mathbf{z}\right) p\left(\boldsymbol{\lambda}\right).$$

The rate posterior can then be determined by the product of the completed likelihood with the gamma prior Eq. (9). For the m'th component of the model this becomes

$$\begin{aligned} p\left(\lambda_m|\boldsymbol{t}, \mathbf{z}\right) &\propto \prod_{n=1}^{\Omega} \lambda_m e^{-\lambda_m t_n} \operatorname{Gamma}\left(\lambda_m; \alpha_m, \beta_m\right) \\ &\propto \prod_{n=1}^{\Omega} \lambda_m e^{-\lambda_m t_n} e^{-\beta_m \lambda_m} \lambda_m^{\alpha_m - 1} \\ &= \lambda_m^{\Omega_m + \alpha_m - 1} e^{-\lambda_m \left(\beta_m + \sum_{\Omega_m} t_n\right)} \end{aligned}$$

where $\Omega_m$ is the number of data points associated with component $m$ of the model, $T_m$ is the sum of event durations for the $\Omega_m$ data points associated with component $m$, and $\alpha_m$ and $\beta_m$ are hyperparameters. After including a normalizing factor, the conditional posterior can be written as a gamma distribution with updated parameters

$$p(\lambda_m|\boldsymbol{t}, \mathbf{z}) = \text{Gamma}(\lambda_m; \alpha_m + \Omega_m, \beta_m + T_m). \tag{S6}$$

**Weights**

For the conditional posterior of the weights ($\boldsymbol{\pi}$), Bayes' theorem reads

$$p(\boldsymbol{\pi}|\boldsymbol{t}, \boldsymbol{\lambda}, \mathbf{z})\, p(\mathbf{z}|\boldsymbol{\lambda}, \boldsymbol{t}) = p(\mathbf{z}|\boldsymbol{\pi}, \boldsymbol{\lambda}, \boldsymbol{t})\, p(\boldsymbol{\pi}|\boldsymbol{\lambda}, \boldsymbol{t}).$$

The posterior for the weights can then be written as

$$p(\boldsymbol{\pi}|\boldsymbol{t}, \boldsymbol{\lambda}, \mathbf{z}) \propto p(\mathbf{z}|\boldsymbol{\pi}, \boldsymbol{\lambda}, \boldsymbol{t})\, p(\boldsymbol{\pi}|\boldsymbol{\lambda}, \boldsymbol{t}).$$

Since the weights are statistically independent of both $\boldsymbol{\lambda}$ and $\boldsymbol{t}$, the equation becomes

$$\begin{aligned} p(\boldsymbol{\pi}|\boldsymbol{t}, \boldsymbol{\lambda}, \mathbf{z}) &\propto p(\mathbf{z}|\boldsymbol{\pi}, \boldsymbol{\lambda}, \boldsymbol{t})\, p(\boldsymbol{\pi}) \\ &\propto \prod_n \pi_{z_n} \lambda_{z_n} e^{-\lambda_{z_n} t_n} \prod_k \pi_k^{\gamma_k - 1} \\ &\propto \prod_k \pi_k^{\Omega_k + \gamma_k - 1}, \end{aligned}$$

where the $\gamma_k$ are hyperparameters. With normalization, the conditional posterior of the weights can be written as a Dirichlet distribution with updated hyperparameters

$$p(\boldsymbol{\pi}|\mathbf{z}) = \text{Dirichlet}_K(\boldsymbol{\pi}; \boldsymbol{\gamma} + \boldsymbol{\Omega}) \tag{S7}$$

where $\Omega_k$ is the number of data points associated with component $k$ of the model and $\boldsymbol{\Omega} = (\Omega_1, \ldots, \Omega_k, \ldots \Omega_K)$.

### S3 Cutoff Comparison

Results for $\beta_2$AR were computed using a cutoff of 6 Å, 7 Å, and 8 Å to determine the sensitivity of results to cutoff (figure S2). The smaller cutoff resulted in shorter binding times and generally smaller $\tau$, while the larger cutoff resulted in larger $\tau$ but generally the same residues were prominent in the $\tau$ vs residue plots. The residues labeled in figure S2A are those residues for which the longest cholesterol binding events have smaller positional variability. All three cutoffs confirm cholesterol binding sites at P88/F89, the cleft between I153 and V164, and at W313.

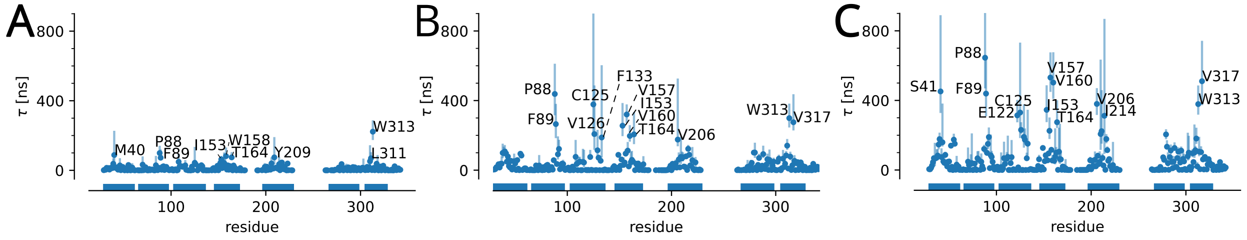


**Figure S2:** $\tau$ vs residue for $\beta_2$AR with cutoff set to 6.0 Å (A), 7.0 Å (B), and 8.0Å (C).

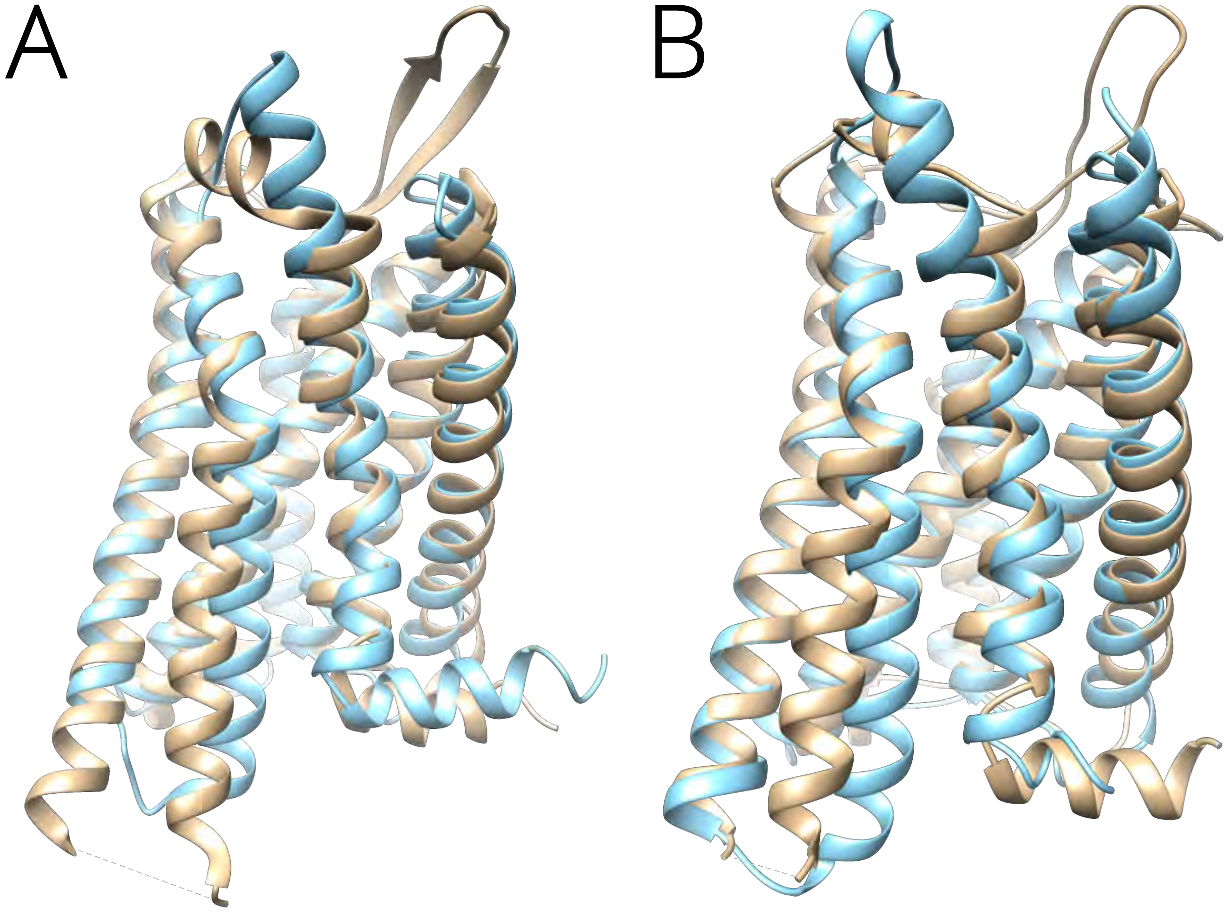


**Figure S3:** CCK model comparison with experimental structures. Comparison of the model (blue) and the experimental structures (tan) for CCK$_1$R (A) and CCK$_2$R (B). Experimental structures depicted are PDB ID 7mbx[6] for CCK$_1$R, and PDB ID 7f8w[7] for CCK$_2$R. Molecular images were rendered with Chimera.[8]

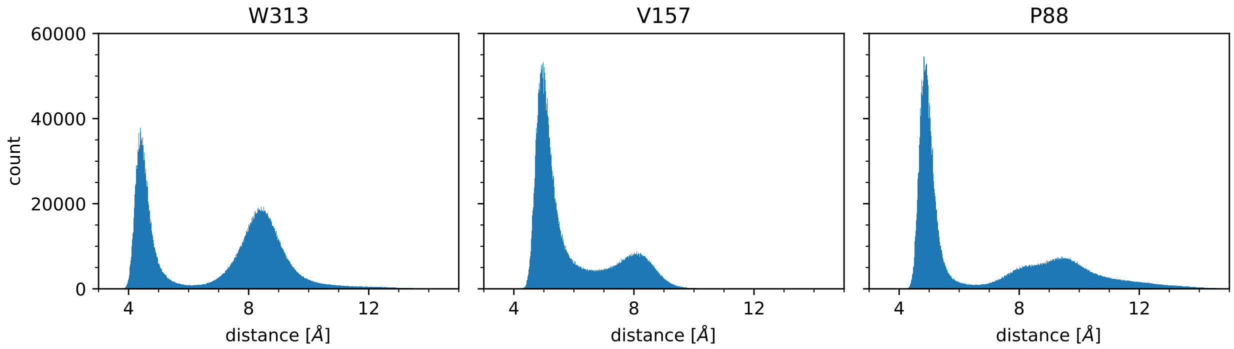


**Figure S4:** Representative protein-cholesterol minimum distance histograms in $\beta_2$AR. The minimum distance between each residue and all cholesterol was computed for each frame and distances were histogrammed. Histograms are shown for (from left to right) W313, V157, and P88.

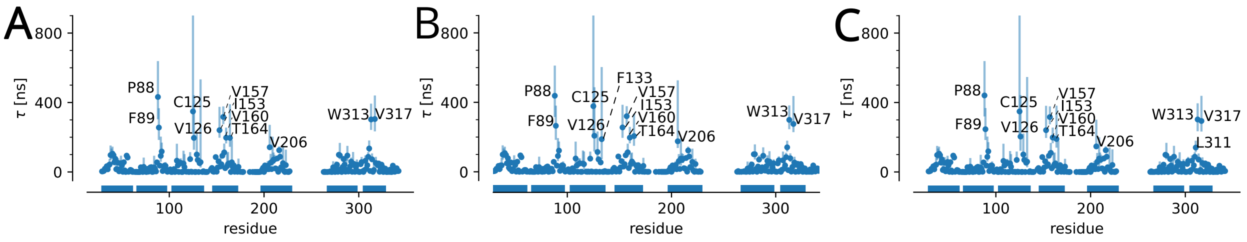


**Figure S5:** $\tau$ vs residue for $\beta_2$AR with burn-in parameter set to (A) 5000, (B) 10000 (as used in the main paper), and (C) 20000.

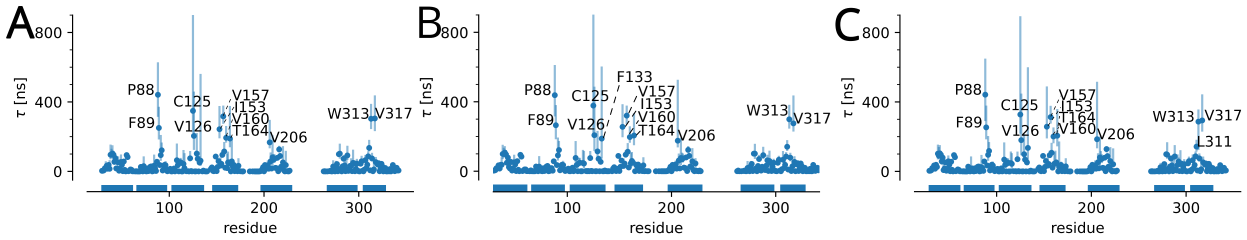


**Figure S6:** $\tau$ vs residue for $\beta_2$AR with thinning parameter set to (A) 50, (B) 100 (as used in the main paper), and (C) 200.

**Table S1:** Rates (in $\text{ns}^{-1}$) and 95%-confidence intervals of clusters in the test data set ($\beta_2$AR W313) for different numbers of components $K$. Because clusters $k \geq 5$ were classified as noise for all $K$, rates corresponding to these clusters are not listed.

| $K$ | 15 | 20 | 50 |
|---|---|---|---|
| $\lambda_0$ | 4.69 [4.66, 4.71] | 4.68 [4.66, 4.71] | 4.68 [4.66, 4.71] |
| $\lambda_1$ | 0.82 [0.80, 0.85] | 0.82 [0.80, 0.85] | 0.83 [0.80, 0.85] |
| $\lambda_2$ | 0.19 [0.16, 0.23] | 0.18 [0.16, 0.23] | 0.19 [0.16, 0.23] |
| $\lambda_3$ | 0.021 [0.017, 0.030] | 0.021 [0.017, 0.028] | 0.021 [0.017, 0.030] |
| $\lambda_4$ | 0.0032 [0.0024, 0.0040] | 0.0033 [0.0025, 0.0040] | 0.0032 [0.0020, 0.0040] |

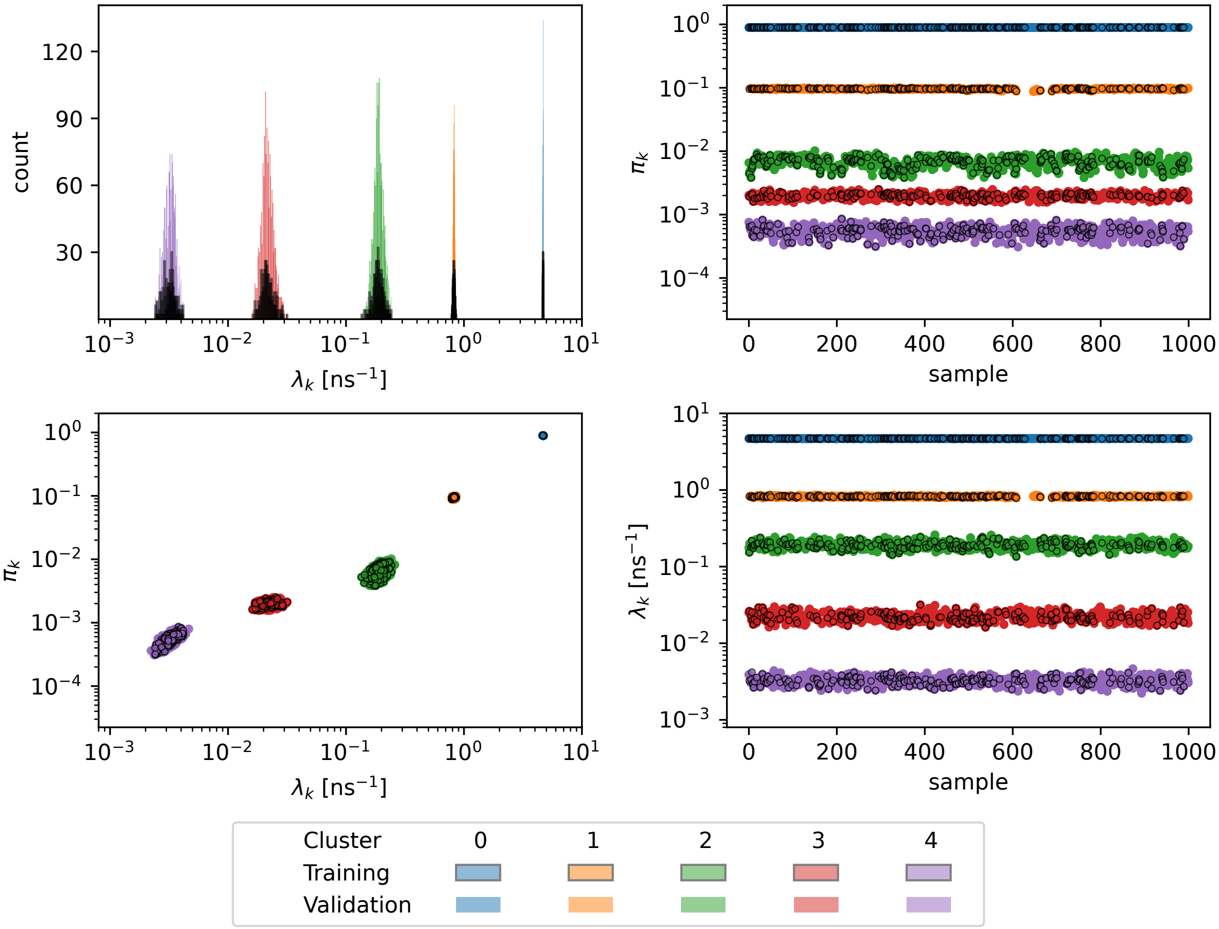


**Figure S7:** Weights and rates for the test data set ($\beta_2$AR W313), using $K = 20$ in the Gibbs sampler. Noise clusters were removed.

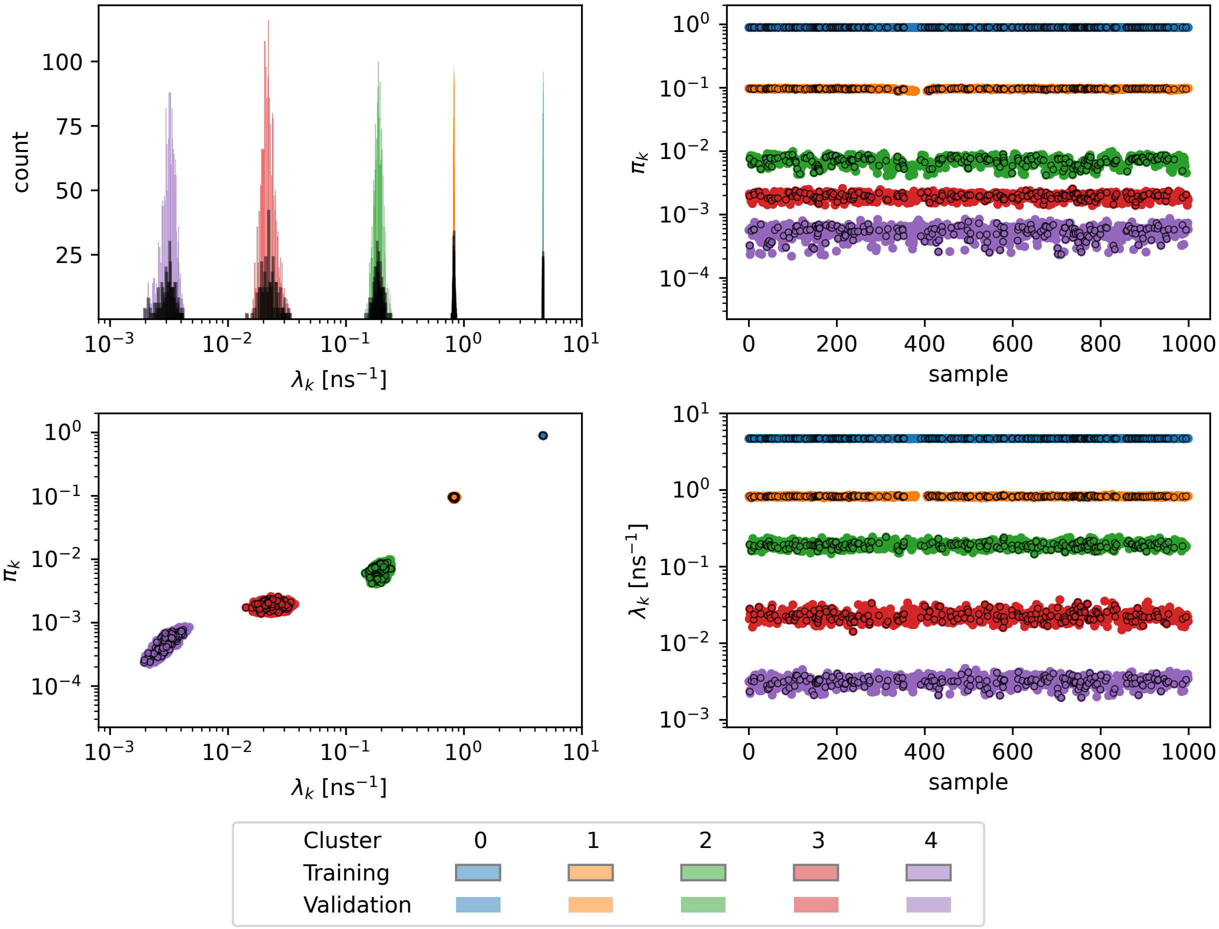


**Figure S8:** Weights and rates for the test data set ($\beta_2$AR W313), using $K = 50$ in the Gibbs sampler. Noise clusters were removed.

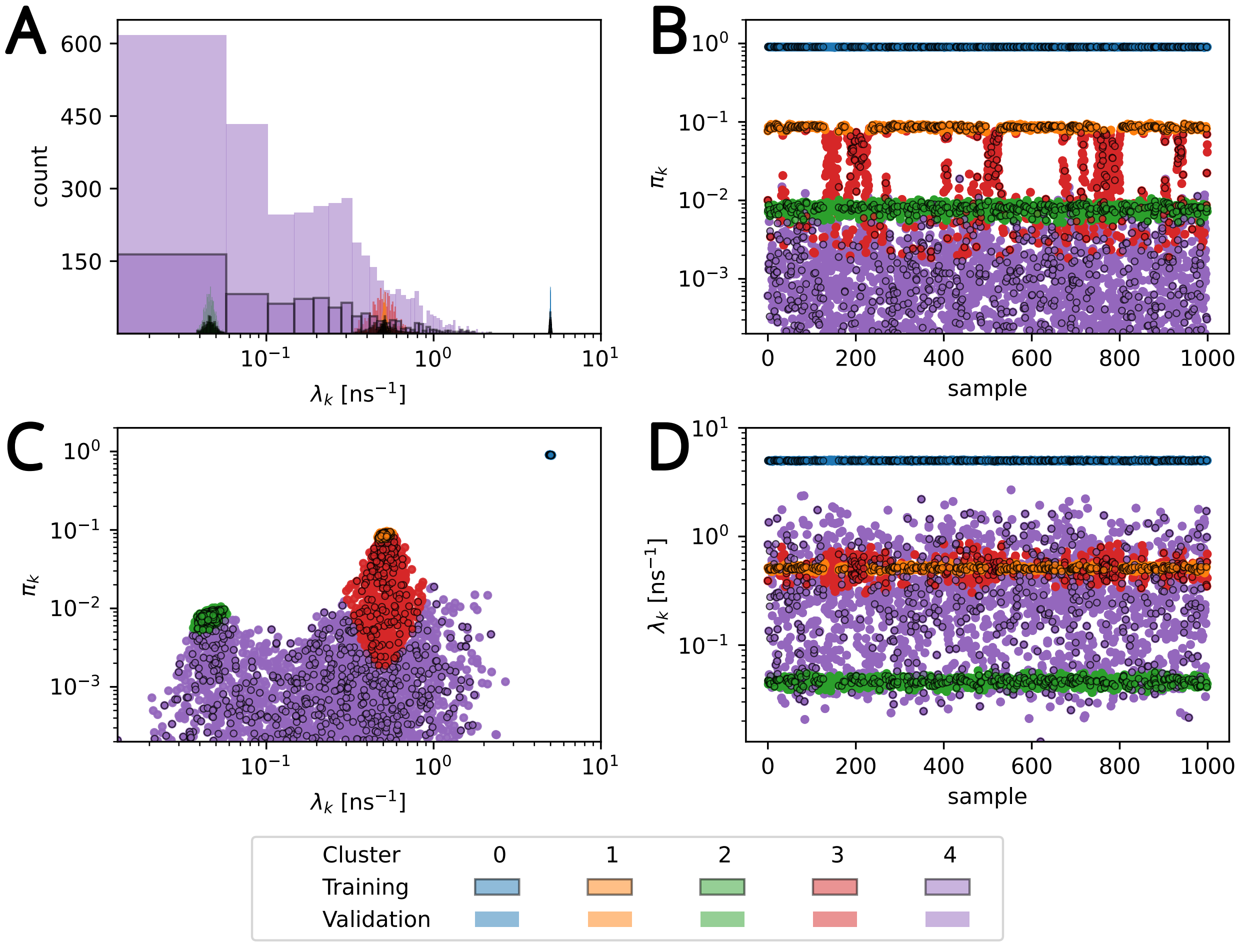


**Figure S9:** Weights and rates for synthetic data. (A) Histograms of rates, (B) weight and (D) rate vs sample, and (C) weight vs rate for thinned Markov chains. Clusters are labeled in order of decreasing rate; samples in the "training" set are used to train the Gaussian mixture model estimator for clustering, and samples labeled "validation" are fit to the trained model.

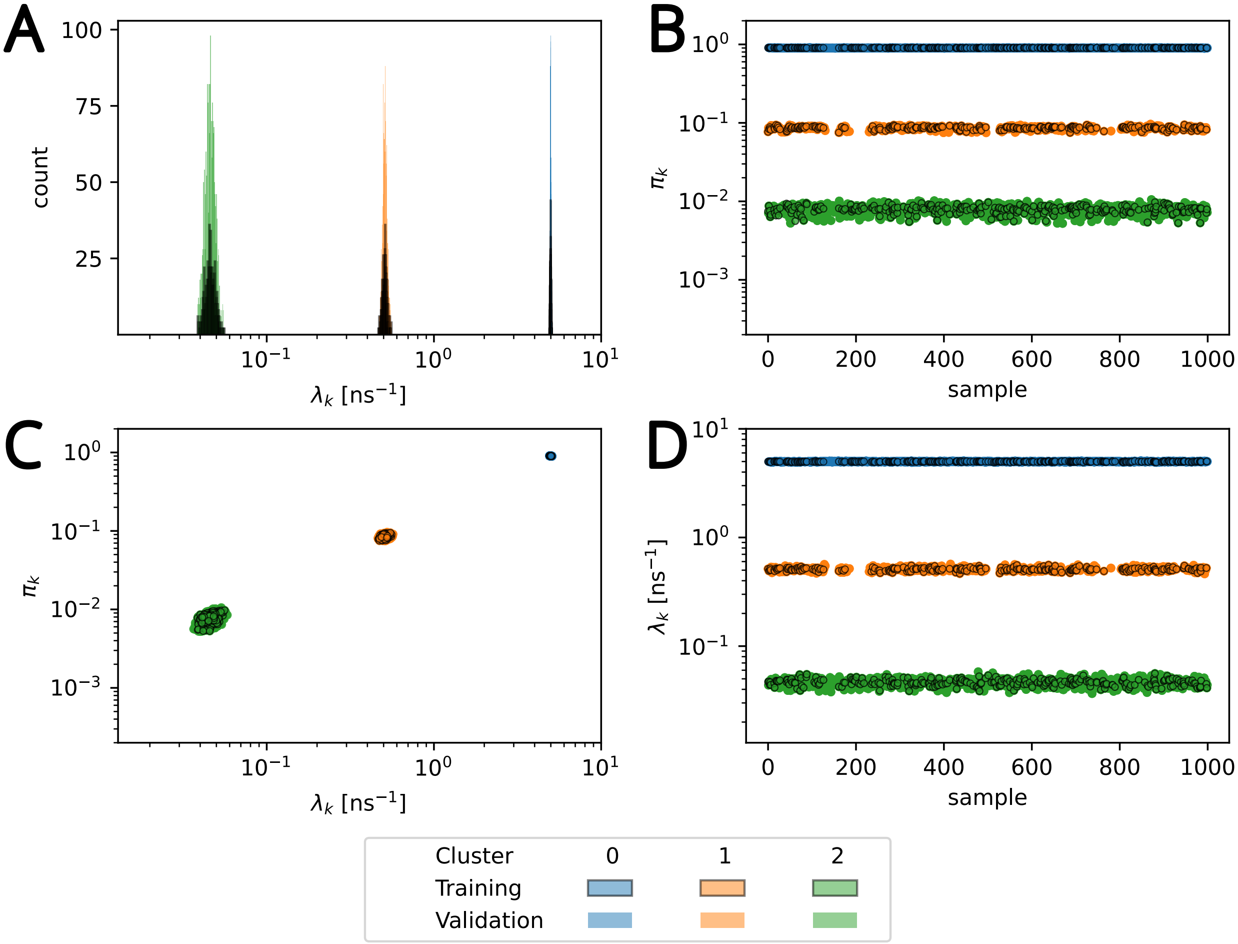


**Figure S10:** Weights and rates for synthetic data with noise clusters removed. (A) Histograms of rates, (B) weight vs sample, (C) weight vs rate, and (D) rate vs sample for thinned Markov chains. Clusters are labeled in order of decreasing rate, "training" samples are used to train the Gaussian mixture model, and "validation" samples are fit to the trained model. Figure S9 contains the results with noise clusters included.

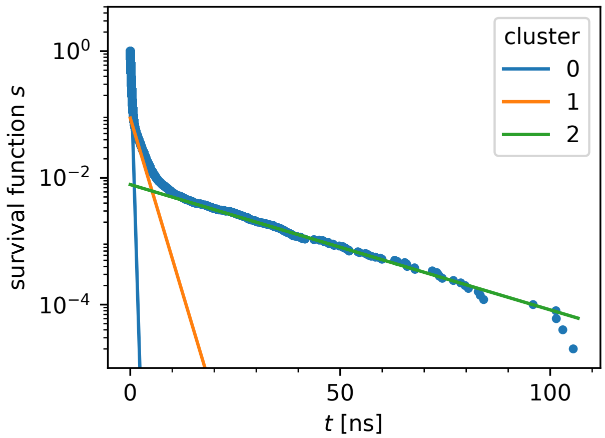


**Figure S11:** Survival function computed from synthetic data with exponential model components, where exponential mixture model parameters were computed from clusters depicted in figure S10.

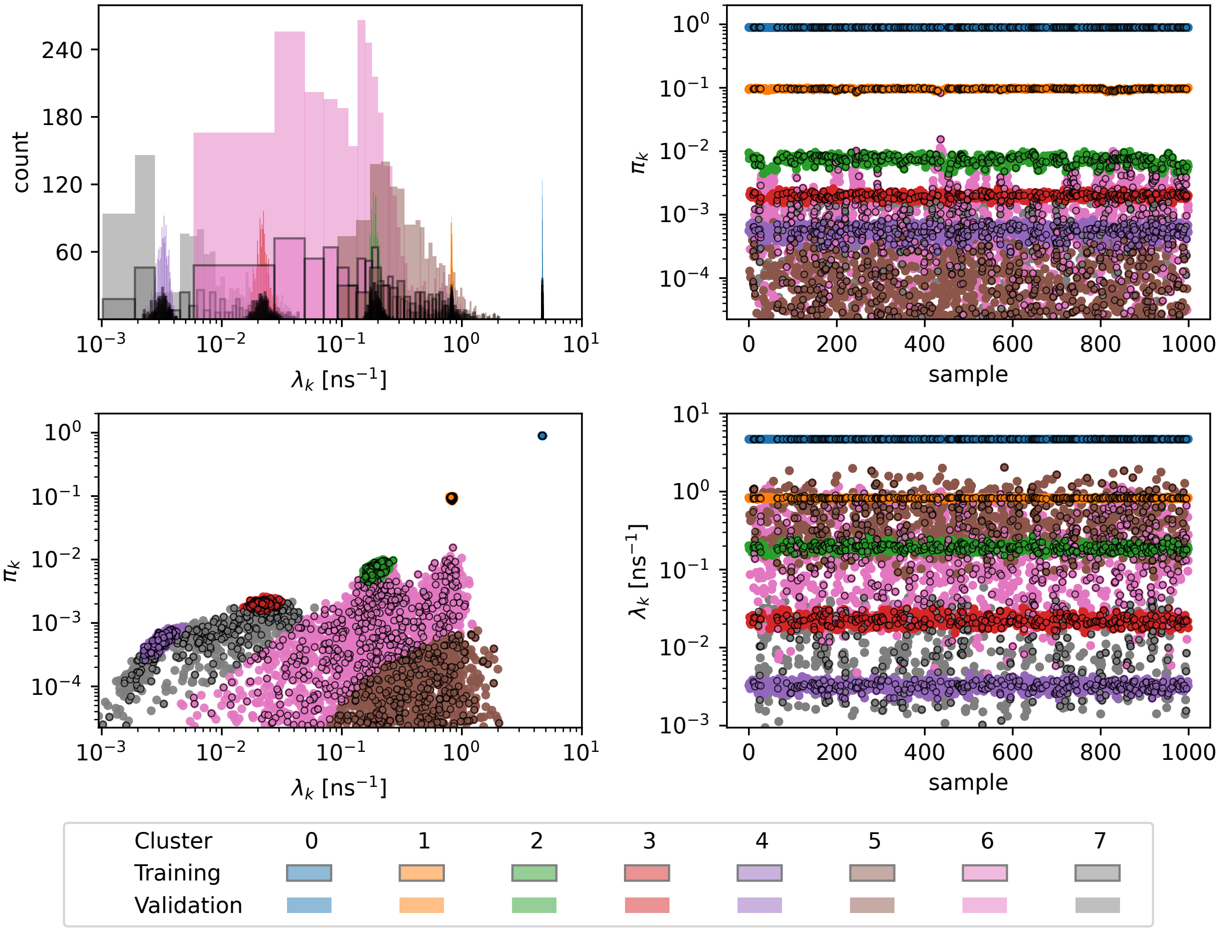


**Figure S12:** Weights and rates for the representative data set ($\beta_2$AR W313). (A) Histograms of rates, (B) weight vs sample, (C) weight vs rate, and (D) rate vs sample for thinned Markov chains. Clusters are labeled in order of decreasing rate, "training" samples are used to train the Gaussian mixture model, and "validation" samples are fit to the trained model. Clusters 5–7 are classified as noise clusters and are removed (see Figure 4 in the main paper).

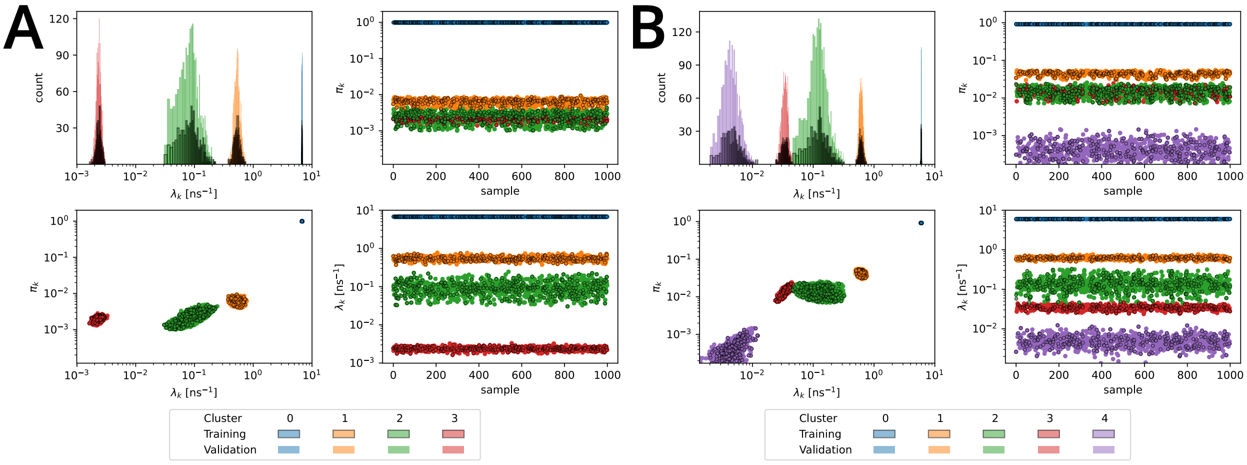


**Figure S13:** Weights and rates for (A) M384 of $CB_1R$ and (B) M286 of $CB_2R$. Noise clusters were removed as described in Methods. Labels have the same meaning as in figure S10.

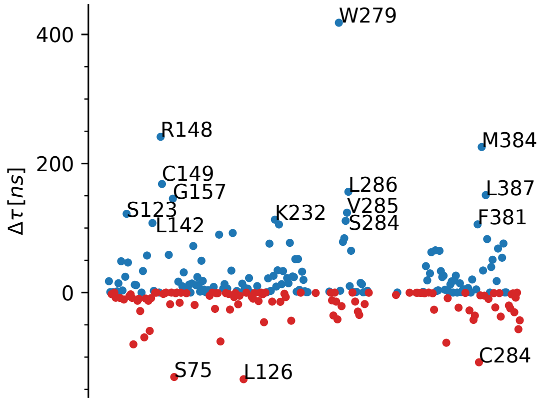


**(S14A)** $CB_1R$ (blue) − $CB_2R$ (red)

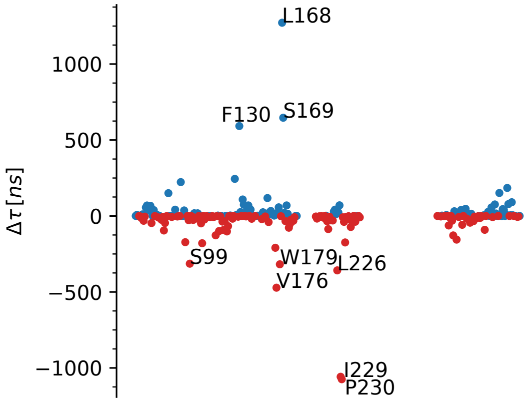


**(S14B)** $CCK_1R$ (blue) − $CCK_2R$ (red)

**Figure S14:** Difference plot for $\tau$ of equivalent residues ($\Delta\tau = \tau_1 - \tau_2$) plotted against residue. Only differences larger than three times the average are labeled with the residue of the protein with the longer $\tau$. Residues with $\Delta\tau > 0$ (blue) belong to either $CB_1R$ or $CCK_1R$ whereas residues with $\Delta\tau < 0$ (red) belong to either $CB_2R$ or $CCK_2R$.

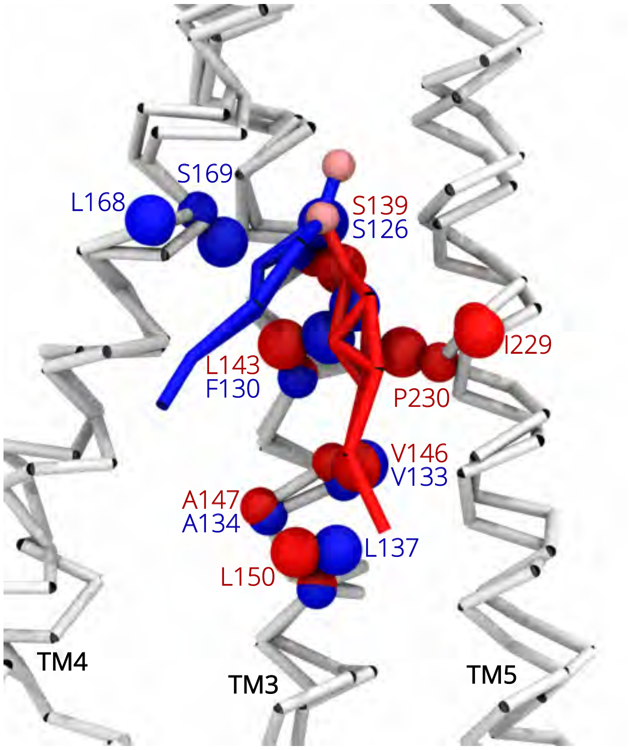


**Figure S15:** Comparison of cholesterol binding sites with long residence times $\tau$ in cholecystokinin receptor homologs $CCK_1R$ (blue) and $CCK_2R$ (red). Residues with long $\tau$ are labeled and shown as CG particles. Cholesterol is shown in representative binding poses, with the OH group particle indicated as a pink sphere. The *kinetic mapping* analysis was performed on equivalent residues, F130 of $CCK_1R$ and L143 of $CCK_2R$. (Image drawn with VMD[9])

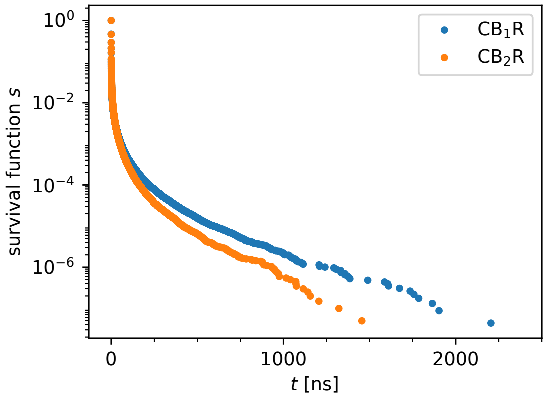


**(S16A)** $CB_1R$ and $CB_2R$

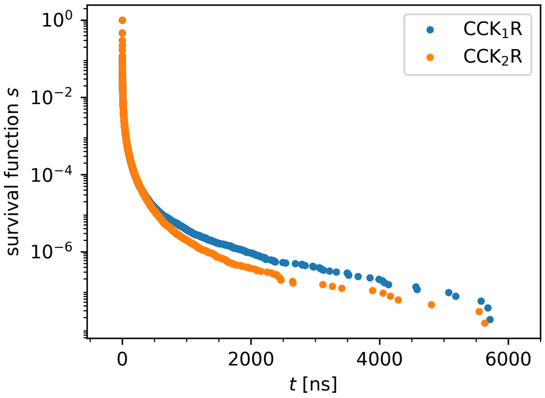


**(S16B)** $CCK_1R$ and $CCK_2R$

**Figure S16:** Whole-protein survival functions of the interaction of cholesterol with any protein residue. The survival function for each protein was computed from the combined contacts from all residues, depicting how the protein overall interacts with cholesterol.

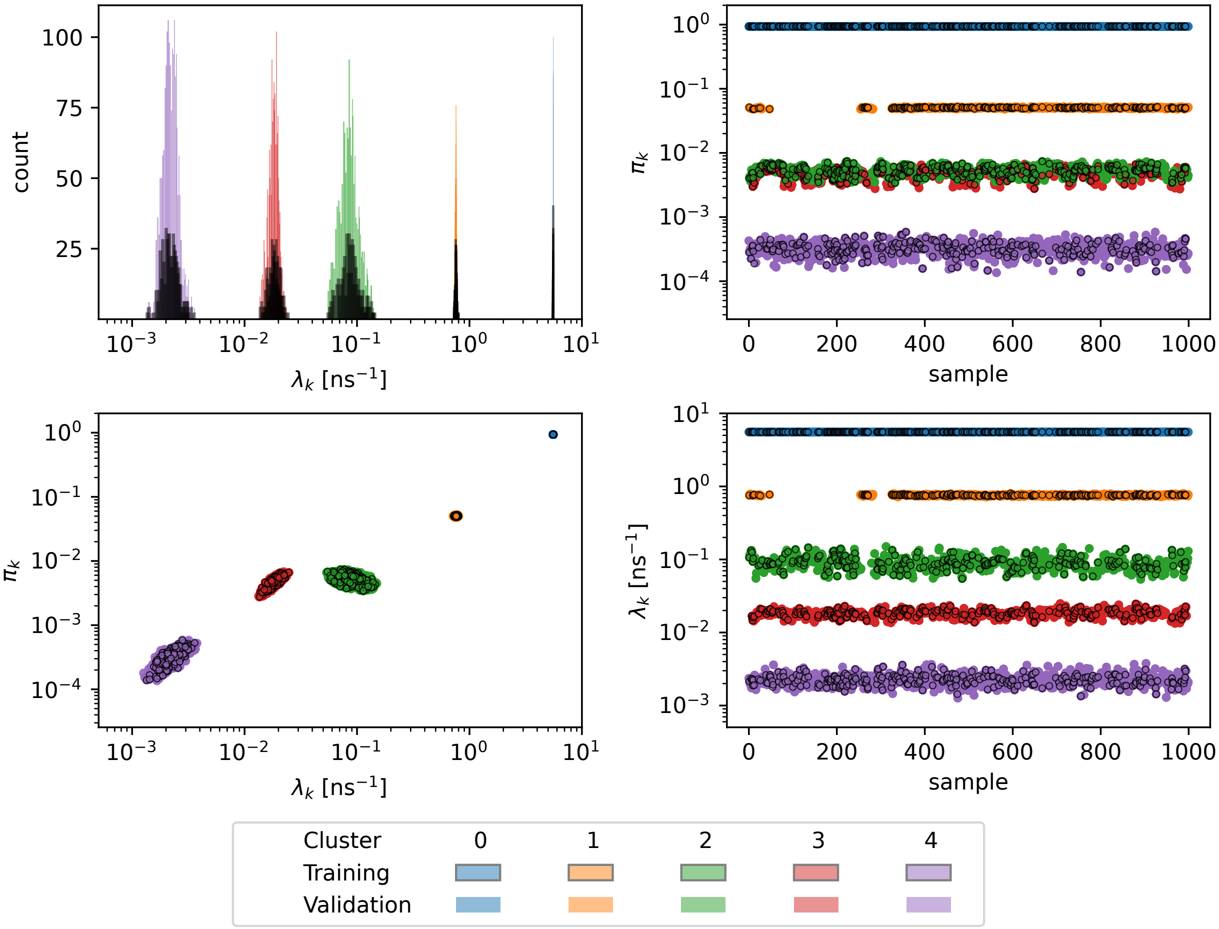


**Figure S17:** Weights and rates for P88 of $\beta_2$AR with noise clusters removed. Labels have the same meaning as in figure S10.

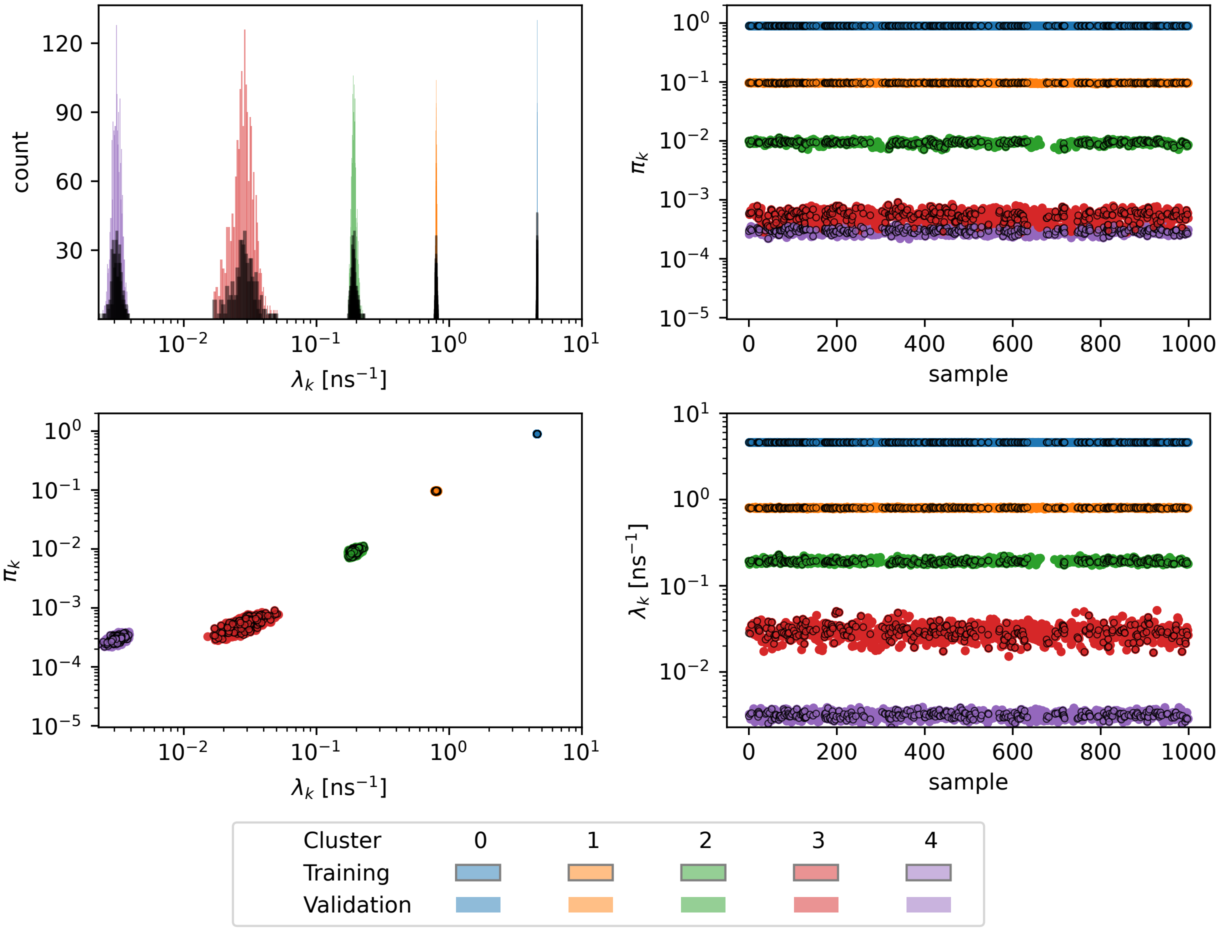


**Figure S18:** Weights and rates for V157 of $\beta_2$AR. with noise clusters removed. Labels have the same meaning as in figure S10.

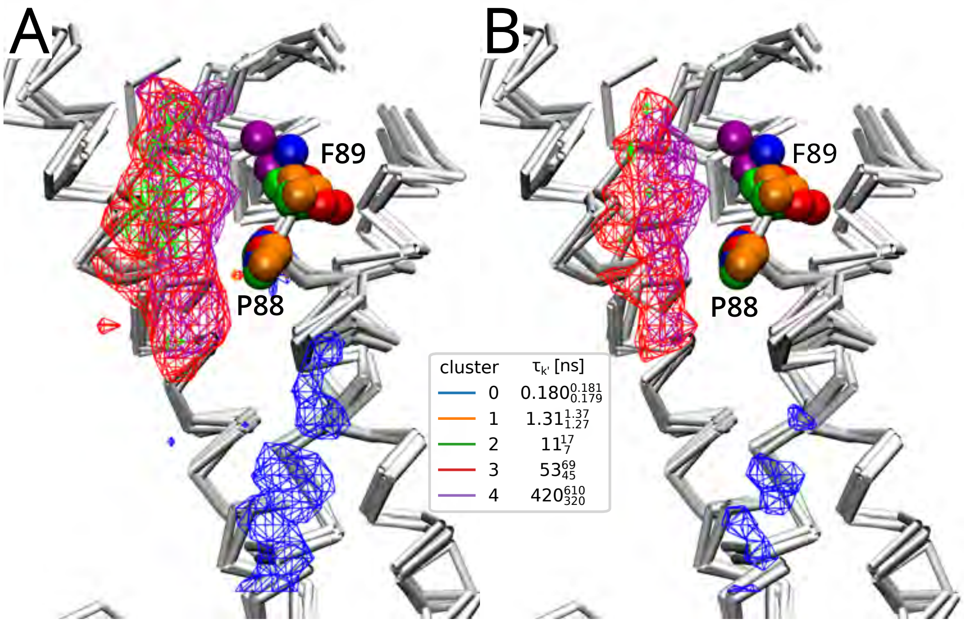


**Figure S19:** Weighted densities for P88 in $\beta_2$AR, where P88 and F89 are shown in the VDW representation and the colors correspond to the P88 equivalent result plot in figure S17. (A) Densities drawn at 0.002 Å$^{-3}$ isocontour cutoff show regions of density near the binding cutoff (7 Å) corresponding to the fastest rate (blue). Several overlapping densities are present due to the three slowest processes (green, red, purple), where densities for the slowest process (purple) are closer to the protein surface and correspond to a contact with the side chain of F89. (B) Densities drawn at 0.05 Å$^{-3}$ show the most dense regions near P88 are due to the slowest two components. The densities and trajectory show the sidechain of F89 pointed toward the cholesterol when the longest binding events are observed. Molecular density images were rendered with VMD.[9]

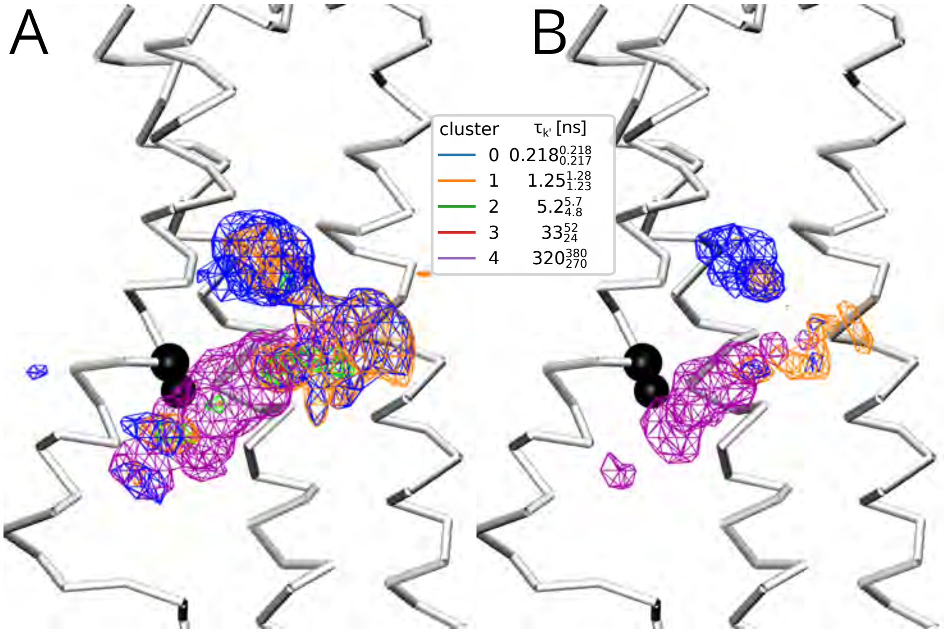


**Figure S20:** Weighted densities for V157 in $\beta_2$AR, where V157 is shown in black. Densities for the fastest two components (blue, orange) are observed in regions near the cutoff where binding times are short. The slowest component (purple) depicts a distinct, elongated region near V157 in which cholesterol is located when the longest binding events occur. Colors correspond to the clustering results shown in figure S18. Densities are shown at isocontour cutoffs (A) 0.002 Å$^{-3}$ (B) 0.005 Å$^{-3}$. Molecular density images were rendered with VMD.[9]

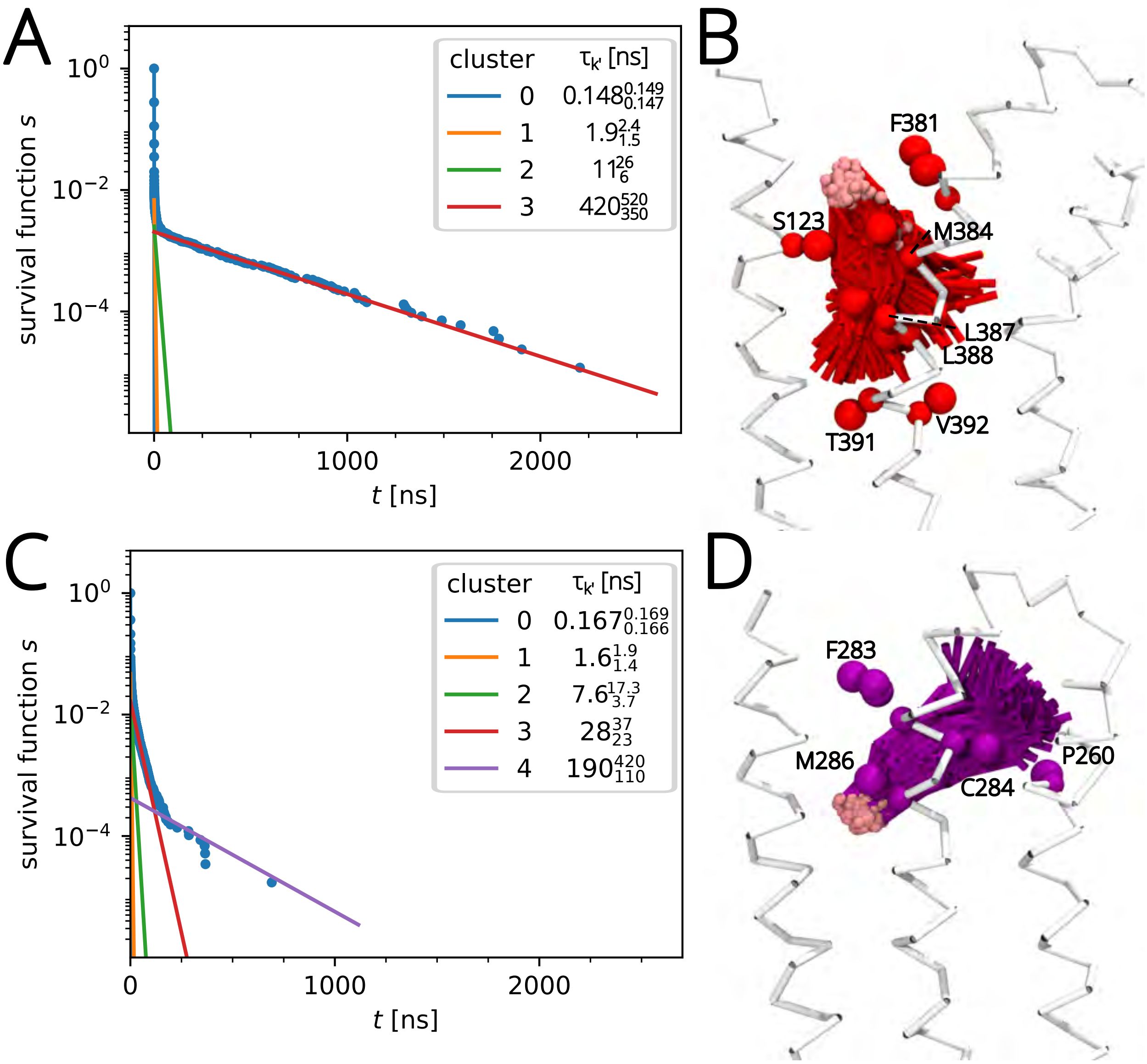


**Figure S21:** Comparison of equivalent sites between cannabinoid receptor homologs. Survival function computed from residence time distribution along with individual components plotted using parameters determined from clustering results for (A) M384 of $CB_1R$ and (C) M286 of $CB_2R$. Molecular images depicting the variability in cholesterol binding poses in the slowest component for (B) M384 of $CB_1R$ and (D) M286 of $CB_2R$, where the pink VDW sphere indicates the polar group in cholesterol. Weights and rates from the Gibbs sampler are included in Figure S13. Molecular images were drawn with VMD.[9]
